# Characterization of a novel glycocin from a thermophile

**DOI:** 10.1101/2025.05.19.655019

**Authors:** Rachel M. Martini, Wilfred A. van der Donk

## Abstract

Glycocins are a growing family of ribosomally synthesized and posttranslationally modified peptides that are *O*– and/or *S-*glycosylated. Using a sequence similarity network of putative glycosyltransferases, the *tht* biosynthetic gene cluster was identified in the genome of *Thermoanaerobacterium thermosaccharolyticum.* ThtA is the precursor peptide to a member of the glycocin F family of glycocins. Like other members of this family, the glycosyltransferase (ThtS) encoded in the biosynthetic gene cluster adds *N-*acetyl-glucosamine to both Ser and Cys residues of ThtA. *S*-linked glycosylation has been shown to be chemically and enzymatically resistant to cleavage and therefore ThtS may be a valuable starting point for engineering efforts. The glycocin derived from ThtA, which we name thermoglycocin, was structurally characterized. Thermoglycocin is unique in that in addition to two nested disulfide bonds, it contains an additional disulfide bond creating a C-terminal loop. Unexpectedly, ThtA lacks the common double glycine motif that denotes a C39-peptidase leader peptide cleavage site. Based on AlphaFold3 modeling, we postulate that cleavage between the leader and core peptide occurs instead at a GK motif. This study adds to the small number of characterized glycocins, employs AlphaFold3 to aid in predicting the structure of the mature peptide product, and suggests a common naming convention similar to that established for lanthipeptides.

**One sentence summary:** Thermoglycocin is a novel glycocin derived from the thermophile *Thermoanaerobacterium thermosaccharolyticum,* containing three disulfide bonds, *O*– and *S*-GlcNAcylation, and is postulated to have a unique C39 protease cut site.

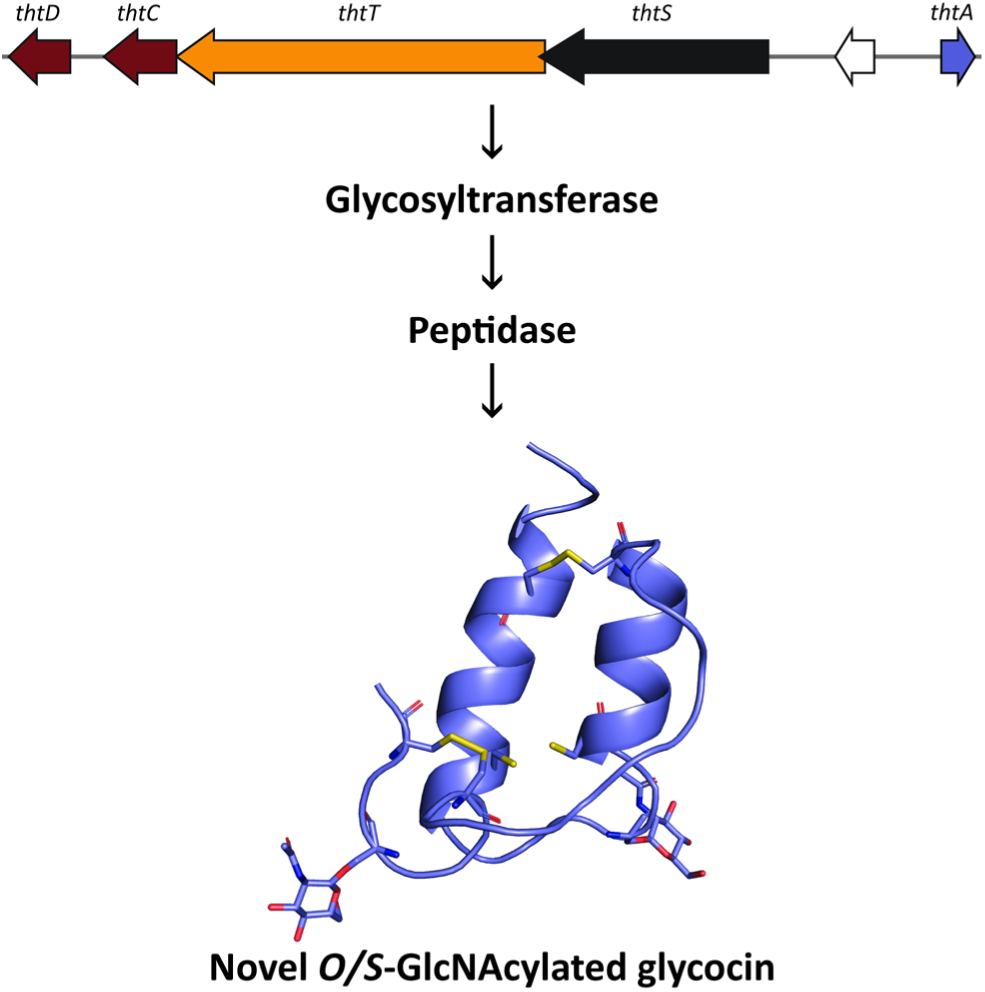

## INTRODUCTION

Recent investigations of ribosomally synthesized and posttranslationally modified peptides (RiPPs) have yielded many novel posttranslational modifications (Nguyen et al., 2024). RiPPs are synthesized as a precursor peptide that contains two regions: a leader peptide and a core peptide (Arnison et al., 2013; Oman & van der Donk, 2010). The core peptide is acted on by modifying enzymes encoded in the biosynthetic gene cluster (BGC), after which the leader peptide is typically cleaved off by proteolysis to create the mature peptide product (Eslami & van der Donk, 2023; Oman & van der Donk, 2010). Glycocins are a growing family of RiPPs, characterized by glycosylations on Ser, Thr, or Cys residues (Hata et al., 2010; Izquierdo et al., 2009; Kaunietis et al., 2019; Main et al., 2020; Maky et al., 2021; Maky et al., 2015; Nagar & Rao, 2017; Norris & Patchett, 2016; Oman et al., 2011; Ren et al., 2018). Compared to *O*-linked glycosylations, *S*-linked glycosylations are rare in biology but they have greater stability and resistance to chemical and enzymatic cleavage (De Leon et al., 2017; Maynard et al., 2016). Previous studies have shown that glycosyltransferases from glycocin biosynthetic pathways can be used to glycosylate non-native substrates (Fujinami et al., 2021; Oman et al., 2011). Thus, the prevalence of *S*-linked glycosylations in glycocins opens up opportunities for application of their glycosyl transferases to other fields (Sharma et al., 2021; Wang et al., 2014). Given the ubiquitous nature of glycosylations with *N-*acetyl-glucosamine (GlcNAc) in eukaryotic organisms (Hart et al., 2011), enzymes that generate *S*-linked GlcNAc modifications are particularly attractive for engineering purposes (Maynard et al., 2016).

In this study, we characterize a novel glycocin in the glycocin F family derived from the thermophilic bacterium *Thermoanaerobacterium thermosaccharolyticum*. A glycocin from a thermophilic strain was chosen for investigation because enzymes from thermophiles generally display high stability that is desirable for engineering and use in biocatalytic processes (Chatterjee et al., 2023; Chettri et al., 2021; Gomes et al., 2016; Zhu et al., 2020). Indeed, other enzymes from this organism have been investigated for their potential use in biotechnology because of their thermostable and robust properties (Pei et al., 2012).

Rapid growth in the field of glycocins suggests that a common naming convention for glycocin biosynthetic machinery may be desirable. A standardized nomenclature using the prefix Lan was proposed more than 30 years ago for the largest class of RiPPs, the lanthipeptides (de Vos et al., 1991), which has proven to be very useful in discussing lanthipeptide families, genes, and enzymes (Repka et al., 2017). Here we propose glycocin precursor peptides to be referred to using the generic term GycA, glycosyltransferases as GycS, peptidases as GycT, and disulfide isomerases as GycC and GycD. Accordingly, the glycocin BGC from *T. thermosaccharolyticum* was given the locus name *tht* and the enzymes encoded in the cluster the names ThtA, ThtS, ThtT, ThtC and ThtD (Fig. 1).

**Fig. 1.**
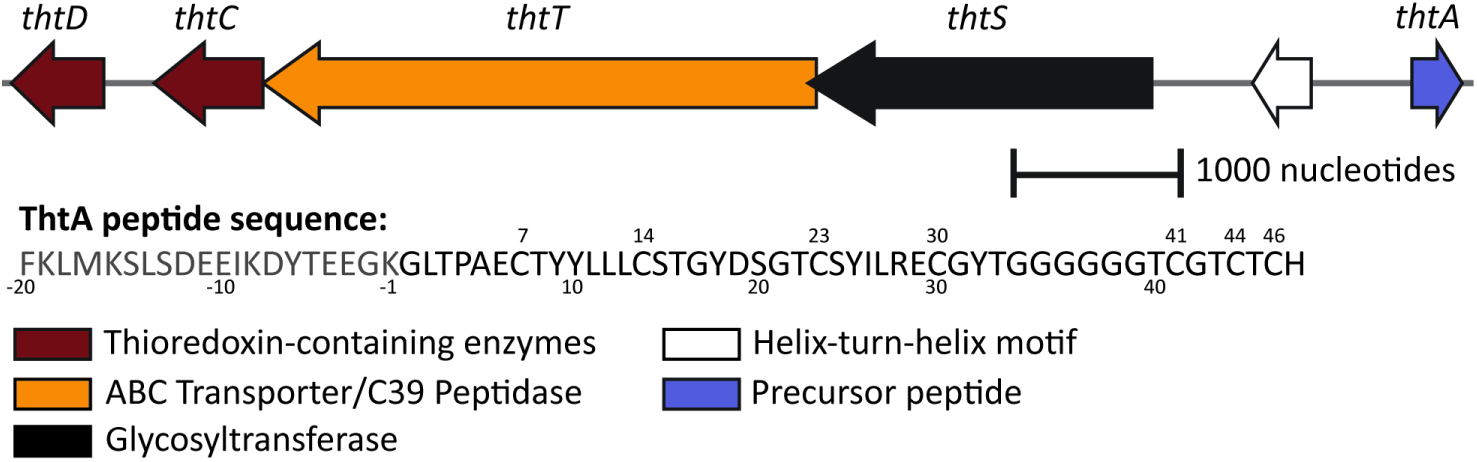
Gene architecture of the *tht* biosynthetic gene cluster and sequence of the precursor peptide ThtA. The end of the leader peptide (residue −1; leader peptide shown in grey) is putative and based on data in this study. Cys residues are numbered above the sequence.

## MATERIALS AND METHODS

### General methods

Chemicals and media for cultures were purchased from Thermo Fisher Scientific or Sigma Aldrich unless otherwise stated. Polymerase chain reactions (PCR) were carried out using a C1000 Bio-Rad thermocycler and were catalyzed using Q5 polymerase (NEB). Matrix-assisted laser desorption/ionization time-of-flight (MALDI-TOF) mass spectrometry analyses were carried out using a Bruker UltrafleXtreme instrument (Bruker Daltonics) through the UIUC Mass Spectrometry facility, using 50 mg/mL super DHB (2,5-dihydroxy benzoic acid) dissolved in 80% acetonitrile and 0.1% trifluoroacetic acid (TFA) as matrix.

### Identification of the *tht* BGC

The sequence similarity network (SSN) (Atkinson et al., 2009) webtool from the enzyme function initiative-enzyme tools (EFI-EST) (Gerlt et al., 2015) was used to generate a sequence similarity network from a UniProt (Consortium, 2024) search of proteins related to glycosyltransferases known to be involved in glycocin biosynthesis. The SSN was visualized using Cytoscape (Shannon et al., 2003) (Supplementary Fig. S1). The network was manually inspected for enzymes that resemble the GlcNAc *S-* glycosyl transferase AsmA (Main et al., 2020) to identify potential orthologs revealing the *tht* BGC.

### Plasmid assembly

Genes encoding *thtA*, *thtS*, and the first 126 residues of *thtT,* containing the C39 domain, were ordered as double stranded DNA from Twist Biosciences. Genes were inserted into pRSFDuet (*thtA* and *thtS*) or pET28b (*thtT*) with HiFi Gibson assembly master mix (NEB). Assembled plasmid was used to transform chemically competent *Escherichia coli* NEB Turbo cells that were then plated on LB agar with the corresponding antibiotic. Plasmid sequence was verified via Sanger DNA sequencing at the UIUC Core Sequencing Facility.

### Heterologous expression of peptides and proteins and *in vitro* modification

Plasmids were used to transform *E. coli* SHuffle cells (NEB) (for co-expression of ThtA and ThtS) or *E. coli* NovaBlue T1^R^ Singles competent cells (for expression of ThtS and MBP-ThtT126). A single colony was used to inoculate small (5 mL) cultures of LB containing 50 mg/L kanamycin; 100 mg/L spectinomycin was added for SHuffle cell cultures. Cultures were grown overnight at 37 °C, while shaking. The small cultures were diluted in 1 L of terrific broth (TB) and grown at 37 °C (30 °C for SHuffle cells) to OD_600_= 0.6. Cultures were incubated on ice for 20 min. Isopropyl ß-D-1-thiogalactopyranoside (IPTG) (GoldBio) was added to 0.5 mM. Cultures were grown overnight at 18 °C (16 °C for SHuffle cells). Cells were harvested via centrifugation at 8000 x*g* for 10 min and resuspended in lysis buffer (20 mM NaH_2_PO_4_ pH 7.5, 500 mM NaCl, 0.5 mM imidazole). Cells were lysed by sonication and cell debris was removed via centrifugation at 45,000 x g for 1 h. Supernatant was loaded onto a Ni-IDA immobilized metal affinity chromatography column (Takara Bio). Beads were washed with 20 mM NaH_2_PO_4_ pH 7.5, 500 mM NaCl, 30 mM imidazole. Peptide or protein was eluted with 20 mM NaH_2_PO_4_ pH 7.5, 100 mM NaCl, 1 M imidazole. The buffer of ThtS was exchanged to protein storage buffer (50 mM HEPES pH 8, 300 mM NaCl, 10% glycerol) using an ultracentrifugal filter (Amicon) with a 30 kDa molecular weight cut off and frozen at –80 °C in small aliquots for later use. The buffer containing purified MBP-tagged ThtT126 was similarly exchanged to 50 mM Tris pH 8.

ThtA was further purified by preparative reversed-phase HPLC (Machery-Nagel C18, 5 μm column) on an Agilent 1260 Infinity II HPLC system. Solvent A contained 0.1% TFA in H_2_O. Solvent B contained 0.1% TFA in acetonitrile. The gradient increased linearly from 2 to 60%B over 35 min.

*In vitro* glycosylation of ThtA by ThtS was performed as described previously for other glycosyl transferases (Wang & van der Donk, 2011) using uridine-5’-diphosphate-α-D-glucose (UDP-Glc) or uridine-5’-diphosphate-α-D-*N*-acetylglucosamine (UDP-GlcNAc; Sigma) under reducing conditions. The reaction was incubated overnight at 25 °C. Peptide was desalted using a C18 SPE column and lyophilized. To facilitate the formation of disulfide bonds in ThtA after glycosylation, the glycosylated peptide was incubated overnight with a mixture of oxidized and reduced glutathione as described previously (Wu et al., 2019).

Fully modified peptide was purified by reversed-phase HPLC using an Agilent 1260 Infinity analytical HPLC with a C18 90 Å 5 µm (Vydac). All mThtA-containing fractions were collected and lyophilized for storage at –20 °C.

### Identification and characterization of post-translational modifications

The identity of the sugar modification was determined by acid-catalyzed hydrolysis and gas chromatography-mass spectrometry (GC-MS) of the cleaved sugar after chemical modification as described previously (Oman et al., 2011), with the exception of a change in standards to *N*-acetylgalactosamine, *N*-acetylglucosamine, and *N*-acetylmannosamine. GC-MS analysis was performed by the Carver Metabolomics Core (University of Illinois Urbana-Champaign Roy J. Carver Biotechnology Center). The number of disulfide bonds was determined by labeling with *N*-ethylmaleimide (NEM) under oxidizing and reducing conditions. Peptide was incubated at 70 °C for 15 min in buffer containing 100 mM sodium citrate (pH 6), 6 M guanidine HCl, 10 mM EDTA, and with or without 10 mM tris(2-carboxyethyl)phosphine (TCEP). The reaction mixture was cooled to room temperature and NEM was added to 10 mM final concentration and the reaction was left in the dark for 30 min at 37 °C. The reaction mixture was desalted with a C18 ZipTip (Agilent) and analyzed by MALDI-TOF MS.

### ThtA structural prediction

To determine the location of GlcNAc modification, ThtA was labeled with NEM as previously described and purified by C18 SPE column. Modified peptide was resuspended in buffer (100 mM Tris pH8, 2 mM CaCl_2_) and digested with chymotrypsin (Worthington Biochemical Corporation) for 1 h at 37 °C. Cleaved peptide was purified by C18 TopTip (Glycen), then analyzed via LC-MS/MS using a C18 column (Kinetex 2.6 μm) and collision-induced dissociation (CID) energy of 30 eV.

To determine the location of the disulfide bonds, ThtA was resuspended in digestion buffer (50 mM Tris pH 8, 0.5 mM CaCl_2_) and digested by thermolysin (Promega) with a 1:100 peptide to enzyme ratio for 1 h at 37 °C. Digested fragments were desalted using C18 TopTip (Glycen) and lyophilized. LC-MS/MS was carried out using a C4 column (Jupiter, 5 μm) and CID energy of 30 eV.

## RESULTS AND DISCUSSION

### Identification of the *tht* biosynthetic gene cluster

To identify potential new GlcNAc *S-*glycosyl transferases, we generated an SSN (Atkinson et al., 2009) using the Enzyme Function Initiative Enzyme Similarity Tool (EFI-EST) version 2024_04/101 (Oberg et al., 2023). We focused on enzymes with similarity to GccB, a glycosyltransferase encoded in the glycocin F gene cluster from *Lactobacillus plantarum* KW30 (Stepper et al., 2011; Venugopal et al., 2011), and AsmA, a glycosyltransferase encoded in the ASM1 gene cluster in *Lactobacillus plantarum* A-1 (Main et al., 2020) (Supplementary Fig. S1). GccB and AsmA both install GlcNAc on Cys residues (Hata et al., 2010; Stepper et al., 2011). The Enzyme Function Initiative genome neighborhood tool (Oberg et al., 2023) was used to identify enzymes encoded in close proximity to the neighbors of GccA and AsmA in the SSN. Since the leader peptides of all known glycocins are removed by a C39-like protease that is part of a bifunctional peptidase-containing ATP-binding transporters (PCAT) (Håvarstein et al., 1995), we focused on glycosyltransferases encoded near such a protein. Furthermore, we used comparison to the sequences of the core peptides of known glycocin precursor peptides to identify systems that were predicted to make glycocins with different structures than previously reported family members. Based on these combined considerations, we chose the *tht* BGC for further study. The ThtA precursor sequence is unique compared to previously characterized glycocins because of the presence of seven Cys residues (Fig. 1), whereas all previously characterized glycocins have five Cys residues (Hata et al., 2010; Izquierdo et al., 2009; Kaunietis et al., 2019; Main et al., 2020; Maky et al., 2021; Maky et al., 2015; Nagar & Rao, 2017; Norris & Patchett, 2016; Oman et al., 2011; Ren et al., 2018). The BGC includes genes encoding a precursor peptide (ThtA), a glycosyltransferase (ThtS), a C39 peptidase/ABC transporter fusion protein (ThtT) (Dirix, Monsieurs, Dombrecht, et al., 2004; Håvarstein et al., 1995), and two thioredoxin-like proteins (ThtCD). The *tht* gene cluster also appears in the compilation of putative Type I glycocin BGCs reported by Singh et al. (Singh & Rao, 2021).

### Characterization of a novel glycocin derived from *T. thermosaccharolyticum*

Co-expression of N-terminally His_6_-tagged ThtA and ThtS in *Escherichia coli* yielded two peptide products showing mass increases of ∼203 Da and ∼402 Da compared to the predicted mass of the unmodified precursor peptide as determined by matrix-assisted laser-desorption/ionization time-of-flight (MALDI-TOF) mass spectrometry (MS) (Supplementary Fig. S2). These mass increases are close to one and two additions of *N*-acetylhexose (expected increase in mass of 203 Da), given the accuracy of the MALDI-TOF mass spectrometer used in this mass range. Indeed, high resolution electrospray ionization (ESI) mass spectrometry verified the addition of two *N*-acetylhexoses (see below). Acid catalyzed hydrolysis was used to cleave the sugars from the peptide followed by their derivatization as described previously (Oman et al., 2011). Gas chromatography monitored by mass spectrometry (GC-MS) and comparison with authentic standards derivatized in the same manner identified the sugar as GlcNAc (Supplementary Fig. S3).

We next turned to *in vitro* characterization of ThtS. His_6_-ThtA and His_6_-ThtS were expressed separately in *E. coli* and purified by nickel-affinity chromatography. *In vitro* reaction of His_6_-ThtA with His_6_-ThtS in the presence of UDP-GlcNAc as a sugar donor resulted in two glycosylations (Fig. 2A) confirming that fully glycosylated ThtA (mThtA) contained two GlcNAc modifications. Use of alternative sugar nucleotides showed that ThtS has some substrate flexibility as the enzyme could also install two glucose molecules (Supplementary Fig. S4), although the reaction is less favored given that mThtA purified from *E. coli*, which contains 0.8-1.5 mM UDP-glucose (Dhamdhere & Zgurskaya, 2010) and ∼ 400 μM UDP-GlcNAc (Namboori & Graham, 2008), was only modified with GlcNAc (Fig. 2A).

**Fig. 2.**
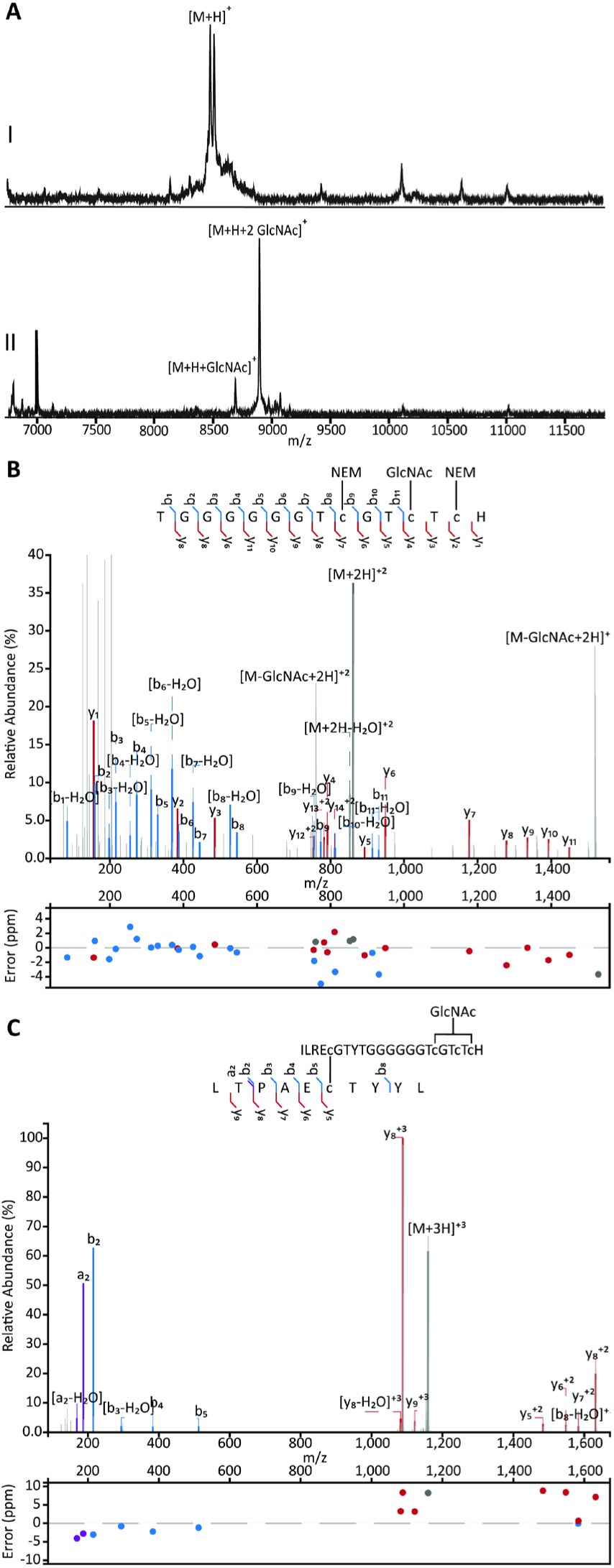
A) MALDI-TOF mass spectra of I) His_6_-ThtA (observed mass: 8507.3, calculated mass: 8507.7), and II) singly (observed mass: 8710.0, calculated mass: 8710.8) and doubly GlcNAcylated (observed mass: 8909.2, calculated mass: 8913.9) mThtA after in vitro incubation with ThtS. B) LC-MS/MS spectrum of residues 33-47 (observed mass: 861.3147, calculated mass: 861.3136) of mThtA GlcNAcylated *in vitro* by ThtS, alkylated with NEM, and digested by chymotrypsin. Fragmentation data is consistent with the +203 modification being on Cys44. Cys41 and Cys46 were alkylated with NEM. C) LC-MS/MS of ThtA residues 2-11 and 26-47 (observed mass: 1159.8359, calculated mass: 1159.8263) connected by a disulfide bond obtained by treating non-reduced mThtA, GlcNAcylated *in vitro* by ThtS, with thermolysin. Panels B and C were prepared using the Interactive Peptide Annotator Webtool (Brademan et al., 2019).

To localize the position of the modified residues, sugar-modified ThtA (mThtA) was reduced with triscarboxyethyl phosphine (TCEP) and the free Cys residues were alkylated with *N-*ethylmaleimide (NEM). The derivatized peptide was digested with chymotrypsin and the fragments analyzed by liquid chromatography tandem mass spectrometry (LC-MS/MS). Fragmentation by collision induced dissociation (CID) showed GlcNAc modification at Cys44 (Fig. 2B) near the C-terminus with Cys41 and Cys46 alkylated by NEM.

The second glycosylation site was more difficult to determine as the second sugar dissociated in the tandem MS experiments, suggesting that the second GlcNAc was *O*-linked, which is a more labile linkage (Drummond et al., 2021; Kaunietis et al., 2019; Main et al., 2020; Stepper et al., 2011). The digestion with chymotrypsin allowed us to narrow down the second glycosylation to a peptide spanning Asp19 through Tyr25 (Supplementary Figure S5). Because of the lability of the GlcNAc during tandem MS analysis, we were not able to conclusively determine whether the site of *O-* glycosylation was Ser20 or Thr22, but based on sequence homology with the glycocins ASM1 and glycocin F (Main et al., 2020; Stepper et al., 2011) the *O-*glycosylation is highly likely to occur on Ser20.

As reported previously for other glycocin glycosyl transferases (Kaunietis et al., 2019; Oman et al., 2011), the leader peptide was not necessary for ThtS activity (Supplementary Fig. S6). Overall, these finding suggest that ThtS is part of a growing class of glycosyltransferases able to catalyze both *O*– and *S*-linked glycosylations (Ahn et al., 2018; Main et al., 2020; Venugopal et al., 2011; Wang et al., 2014).

### Determination of the disulfide pattern

All glycocins characterized thus far have disulfide bonds that contribute to their remarkable stability and that play an important role in glycocin bioactivity (Bisset et al., 2018; Dorenbos et al., 2002). Reaction of mThtA isolated from *E. coli* with *N*-ethylmaleimide (NEM) in the absence of reductant did not lead to any alkylation suggesting the peptide does not contain any free cysteine residues (Supplementary Fig. S7). Repeating the reaction in the presence of TCEP resulted in the addition of six NEM molecules (Supplementary Fig. S8), suggesting all six Cys residues that are not glycosylated are involved in disulfide bonding. All well-characterized glycocins to date have two nested disulfide bonds, connecting two α-helices (Garcia De Gonzalo et al., 2014; Venugopal et al., 2011). Previous studies have employed PSIPRED workbench (Buchan & Jones, 2019) to aid in assigning secondary structure to glycocins (Kaunietis et al., 2019; Wang et al., 2014). Because of the recent advancements in structure prediction technology, AlphaFold3 (Abramson et al., 2024) was used in the current study along with PSIPRED to create a model of ThtA and precursor peptides of known glycocins to gauge the accuracy of the models. PSIPRED (Supplementary Fig. S9) and AlphaFold3 (Supplementary Fig. S10) both predict ThtA to contain two α-helices connected by a flexible linker, and a C-terminal random coil. AlphaFold3 also predicted two nested disulfide bonds between residues Cys7 and Cys30, and between Cys14 and Cys23, similar to the experimentally determined disulfide bond pattern in sublancin (Garcia De Gonzalo et al., 2014; Oman et al., 2011) and glycocin F (Venugopal et al., 2011). AlphaFold3 predicts the third disulfide bond to be between Cys41 and Cys46 to create a C-terminal loop (Fig. 3A). The per-residue measure of local confidence (pLDDT) values are generally very high (>70) in the two helices, with lower confidence in the N– and C-terminal regions and the loop connecting the two helices (Supplementary Fig. S10).

**Fig. 3.**
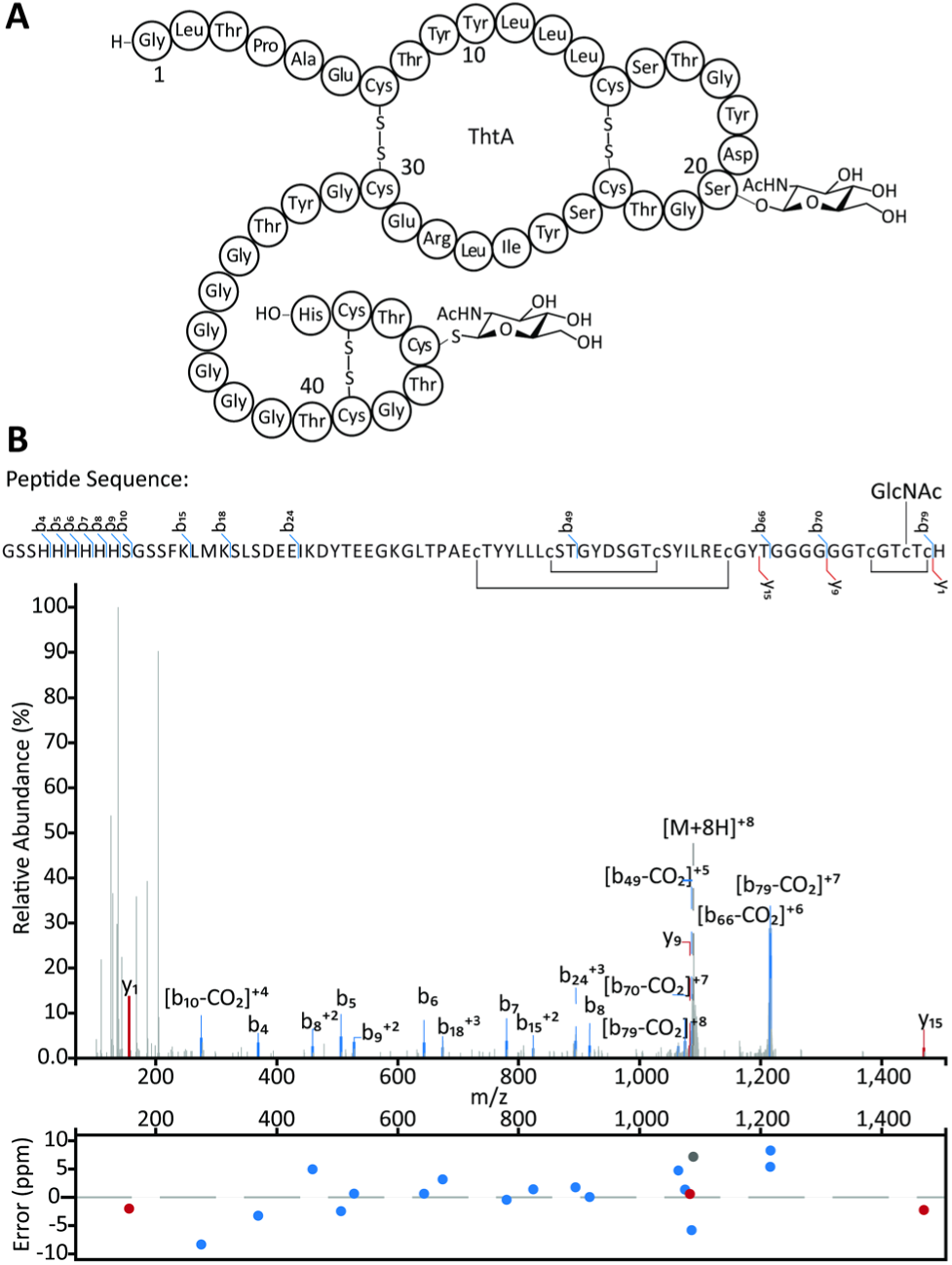
Structure determination of the glycocin from *T. thermosaccharolyticum.* (A) Predicted structure of thermoglycocin based on the AlphaFold3 model, proteolytic digests, and tandem MS analysis. (B) LC-MS/MS data of ThtA (observed mass: 1088.9804, calculated mass: 1088.9726) expressed in *E. coli* carrying one GlcNAc molecule. Panel B prepared using the Interactive Peptide Annotator Webtool (Brademan et al., 2019).

To experimentally confirm this model, mThtA was cleaved with thermolysin in the absence of reductant. Fragments were analyzed using LC-MS/MS. A triply charged ion of 1159.8263 Da was observed that corresponds to the mass of residues 2 to 11 of ThtA connected via a disulfide bond between Cys7 and Cys30 to a peptide consisting of residues 26 to 47 (Fig. 2C). Moreover, an observed singly charged ion of 1467.5767 Da corresponds to residues 12-25 with a disulfide bond between Cys14 and Cys23 (Supplementary Fig. S11). The full length mThtA peptide was also analyzed by LC-MS/MS, showing minimal fragmentation between residues Lys−8 and Tyr33 and no fragmentation between Cys41 and Cys46 (Fig. 3B). As no or little fragment ions are expected within a ring structure, these observations support the proposed disulfide bonding pattern predicted by AlphaFold3 (Fig. 3A). These data suggest therefore that AlphaFold3 may be useful in predicting the folding of other glycocins in future work.

### Prediction of the leader peptide cleavage site

Almost all RiPPs are made from a precursor peptide that contains an N-terminal leader peptide that is removed in a late biosynthetic step by a protease (Eslami & van der Donk, 2023; Montalbán-López et al., 2021). The responsible protease is often encoded within the BGC, and indeed the *tht* BGC encodes an ATP-dependent transporter with a C39-type N-terminal protease (ThtT). These bifunctional peptidase-containing ATP-binding transporters (PCATs) are frequently found in RiPP BGCs (Ayikpoe et al., 2022; Eslami & van der Donk, 2023; Håvarstein et al., 1995). Most characterized C39 peptidases have been shown to cleave after GG, GA, or GS motifs, typically called the double glycine motif (Bobeica et al., 2019; Dirix, Monsieurs, Dombrecht, et al., 2004; Dirix, Monsieurs, Marchal, et al., 2004; Ishii et al., 2010). Given the lack of such a motif in ThtA (Fig. 1), it is not clear what the leader peptide removal site is in ThtA. We made a multiple sequence alignment with related peptides retrieved from the NCBI database using BLAST (Boratyn et al., 2013) (Fig. 4). The structure of glycocin F (Stepper et al., 2011; Venugopal et al., 2011) suggests that its precursor peptide GccF is cleaved at a double glycine motif. The double Gly motif of GccF (GG-K) corresponds to the sequence GK-G in ThtA, which was also hypothesized to be the leader peptide removal site in the glycocin Hyp1 (Fig. 4) (Kaunietis et al., 2019). However, this predicted proteolytic site is unusual for a number of reasons. First, previously studied C39 peptidases that are part of the PCAT family were shown to be unable to accept charged residues in the −1 position (Furgerson Ihnken et al., 2008). Second, most PCAT substrates have hydrophobic residues (Leu, Val, Ile, Met) in positions −7 and −12, which occupy hydrophobic pockets in the PCAT as shown by X-ray crystallography (Bobeica et al., 2019). An example is the precursor peptide sequence to the glycocin sublancin (Fig. 4). But for all the precursor peptides other than sublancin shown in Fig. 4, the residues at position −7 are hydrophilic (Asp/Glu/Ser/Thr/Asn/Lys) and at position −12 mostly negatively charged (Glu/Asp). We noted that the residues at positions −9 and −14 in these peptides are invariable hydrophobic (Leu, Ile, Val, Fig. 4), possibly suggesting that the register has shifted in this set of peptides. To investigate this hypothesis we used AlphaFold3 modelling of the ThtA leader peptide and the protease domain of ThtT encompassing the N-terminal 126 residues (ThtT126, Supplementary Fig. S12). The model showed that residues Ile−9 and Leu−14 fit into a hydrophobic groove on ThtT126 and that Lys−1 aligns with the catalytic Cys in the active site of the peptidase (Bobeica et al., 2019; Chen et al., 2001; Ishii et al., 2010). Thus, it seems that the group of PCATs that cleave substrates with a GKG sequence are able to do so by setting a different register in which now hydrophobic residues at positions −9 and −14 occupy the same pockets that normally are occupied by residues at positions −7 and −12. The known cleavage site to release glycocin F provides indirect support for such a change in register. The precursor peptide for this glycocin has the canonical double Gly motif, but it also has the hydrophobic residues in positions −9 and −14 (Fig. 4).

**Fig. 4.**
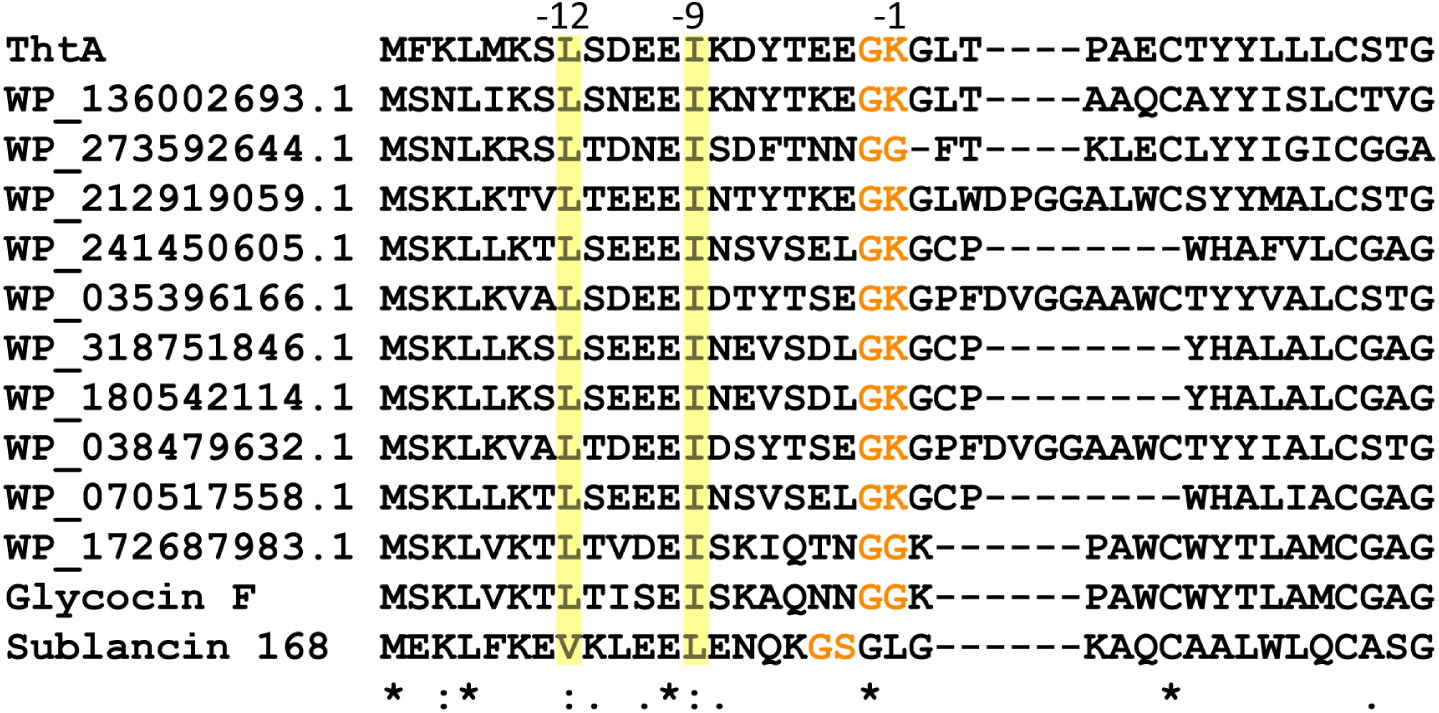
Multiple sequence alignment of ThtA with the leader peptide region of precursor peptides of related putative and known glycocins.

In an attempt to experimentally verify these predictions, the N-terminal C39 protease domain of ThtT was expressed in *E. coli* as has been done previously for the orthologous enzymes CvaB (Wu & Tai, 2004), ComA (Ishii et al., 2006), LahT (Bobeica et al., 2019), LctT (Furgerson Ihnken et al., 2008), BovT (Wang et al., 2016), ColT (Wang et al., 2025), McyT (H. Wang et al., 2023), and MfuT (X. Wang et al., 2023). Unfortunately, both a His_6_-tagged version and a maltose binding protein fusion protein were obtained in insoluble form under a variety of expression conditions. Because of the inability to reconstitute the activity of the C39 protease domain of ThtT, endoproteinase LysC was used with mThtA to produce the putative final compound. Antimicrobial assays against a panel of bacteria did not show any growth inhibition activity (Table S1), which may either indicate that the predicted leader peptide removal site is not correct or that the activity is narrow spectrum and sensitive bacteria were not in the panel investigated. Further studies will be needed to address this question.

The structure in Fig. 3A and Supplementary Fig. S10 suggests a similar topology for the glycocin from *T. thermosaccharolyticum* as previously characterized compounds like sublancin and glycocin F. Furthermore, the model predicts Arg28 to be in a similar location to that of Arg33 in sublancin 168. The presence of a basic residue in this position was shown to be important for the antimicrobial activity of sublancin 168 (Biswas et al., 2017). Basic residues have been predicted in similar locations on Hyp1, Hyp2, pallidocin, and thurandacin A (Kaunietis et al., 2019; Maky et al., 2015; Ren et al., 2018). The conservation of these positively charged residues suggests these glycocins may have a similar target. Because their antimicrobial activity can be antagonized by free sugars, the glycocins for which the mode of action has been investigated are thought to exert their bioactivities with the involvement of phosphoenolpyruvate:sugar phospho-transferase systems (PTSs) such as the glucose (Biswas et al., 2017; Garcia De Gonzalo et al., 2015; Wu et al., 2019) or GlcNAc (Bisset et al., 2018; Norris & Patchett, 2016) transporters. Although we were unable to detect growth inhibitory activity with the tested organisms, given the conserved features in the glycocin from *T. thermosaccharolyticum,* it is tempting to speculate that its activity will also involve the GlcNAc PTS.

## Conclusions

Thermoglycocin is the first glycocin derived from an anaerobic thermophile to be characterized. It is a member of a growing group of *S*-linked glycopeptides, joining the well-characterized glycocins sublancin 168, thurandacin, and glycocin F, a group of peptides that contain *S*-linked GlcNAc modifications. As others have predicted, the diversity of glycocin structures will continue to grow as more are discovered (Palaniappan et al., 2020; Singh & Rao, 2021). As shown in other studies on small, disulfide-containing proteins (Gai et al., 2025; Pan et al., 2025), AlphaFold3 appears to be a useful tool to predict disulfide bonding patterns for glycocins that have additional Cys residues beyond the canonical four Cys residues that form a nested disulfide.

## Acknowledgements

The authors thank the metabolomics facility of the Roy J. Carver Biotechnology Center (CBC) at the University of Illinois at Urbana-Champaign for GC-MS and high resolution mass spectrometry services, and Enleyona Weir (UIUC) for help with the antimicrobial activity assays. HHMI lab heads have previously granted a nonexclusive CC BY 4.0 license to the public and a sublicensable license to HHMI in their research articles. Pursuant to those licenses, the author-accepted manuscript of this article can be made freely available under a CC BY 4.0 license immediately upon publication.

## Supplementary Material

Supplementary material is available online at JIMB (www.academic.oup.com/jimb).

## Author Contributions

R.M.M. and W.A.v.d.D. designed the study. R.M.M. performed all experiments. R.M.M. and W.A.v.d.D. analyzed the data, and R.M.M. and W.A.v.d.D. wrote the manuscript.

## Data Availability

All data are incorporated into the article and its online Supplementary Material. Primary data are deposited at Mendeley Data:

## References

1. Abramson, J., Adler, J., Dunger, J., Evans, R., Green, T., Pritzel, A., Ronneberger, O., Willmore, L., Ballard, A. J., Bambrick, J., Bodenstein, S. W., Evans, D. A., Hung, C.-C., O’Neill, M., Reiman, D., Tunyasuvunakool, K., Wu, Z., Žemgulytė, A., Arvaniti, E., Beattie, C., Bertolli, O., Bridgland, A., Cherepanov, A., Congreve, M., Cowen-Rivers, A. I., Cowie, A., Figurnov, M., Fuchs, F. B., Gladman, H., Jain, R., Khan, Y. A., Low, C. M. R., Perlin, K., Potapenko, A., Savy, P., Singh, S., Stecula, A., Thillaisundaram, A., Tong, C., Yakneen, S., Zhong, E. D., Zielinski, M., Žídek, A., Bapst, V., Kohli, P., Jaderberg, M., Hassabis, D., & Jumper, J. M. (2024). Accurate structure prediction of biomolecular interactions with AlphaFold 3. Nature, 630(8016), 493–500.

2. Ahn, S., Stepper, J., Loo, T. S., Bisset, S. W., Patchett, M. L., & Norris, G. E. (2018). Expression of *Lactobacillus plantarum* KW30 *gcc* genes correlates with the production of glycocin F in late log phase. FEMS Microbiol. Lett., 365(23).

3. Arnison, P. G., Bibb, M. J., Bierbaum, G., Bowers, A. A., Bugni, T. S., Bulaj, G., Camarero, J. A., Campopiano, D. J., Challis, G. L., Clardy, J., Cotter, P. D., Craik, D. J., Dawson, M., Dittmann, E., Donadio, S., Dorrestein, P. C., Entian, K. D., Fischbach, M. A., Garavelli, J. S., Göransson, U., Gruber, C. W., Haft, D. H., Hemscheidt, T. K., Hertweck, C., Hill, C., Horswill, A. R., Jaspars, M., Kelly, W. L., Klinman, J. P., Kuipers, O. P., Link, A. J., Liu, W., Marahiel, M. A., Mitchell, D. A., Moll, G. N., Moore, B. S., Müller, R., Nair, S. K., Nes, I. F., Norris, G. E., Olivera, B. M., Onaka, H., Patchett, M. L., Piel, J., Reaney, M. J., Rebuffat, S., Ross, R. P., Sahl, H. G., Schmidt, E. W., Selsted, M. E., Severinov, K., Shen, B., Sivonen, K., Smith, L., Stein, T., Süssmuth, R. D., Tagg, J. R., Tang, G. L., Truman, A. W., Vederas, J. C., Walsh, C. T., Walton, J. D., Wenzel, S. C., Willey, J. M., & van der Donk, W. A. (2013). Ribosomally synthesized and post-translationally modified peptide natural products: overview and recommendations for a universal nomenclature. Nat. Prod. Rep., 30(1), 108–160.

4. Atkinson, H. J., Morris, J. H., Ferrin, T. E., & Babbitt, P. C. (2009). Using sequence similarity networks for visualization of relationships across diverse protein superfamilies. PLoS One, 4(2), e4345.

5. Ayikpoe, R. S., Shi, C., Battiste, A. J., Eslami, S. M., Ramesh, S., Simon, M. A., Bothwell, I. R., Lee, H., Rice, A. J., Ren, H., Tian, Q., Harris, L. A., Sarksian, R., Zhu, L., Frerk, A. M., Precord, T. W., van der Donk, W. A., Mitchell, D. A., & Zhao, H. (2022). A scalable platform to discover antimicrobials of ribosomal origin. Nat. Commun., 13(1), 6135.

6. Bisset, S. W., Yang, S. H., Amso, Z., Harris, P. W. R., Patchett, M. L., Brimble, M. A., & Norris, G. E. (2018). Using chemical synthesis to probe structure-activity relationships of the glycoactive bacteriocin glycocin F. ACS Chem. Biol., 13(5), 1270–1278.

7. Biswas, S., Garcia De Gonzalo, C. V., Repka, L. M., & van der Donk, W. A. (2017). Structure-activity relationships of the S-linked glycocin sublancin. ACS Chem. Biol., 12(12), 2965–2969.

8. Bobeica, S. C., Dong, S. H., Huo, L., Mazo, N., McLaughlin, M. I., Jimenéz-Osés, G., Nair, S. K., & van der Donk, W. A. (2019). Insights into AMS/PCAT transporters from biochemical and structural characterization of a double glycine motif protease. eLife, 8, e42305.

9. Boratyn, G. M., Camacho, C., Cooper, P. S., Coulouris, G., Fong, A., Ma, N., Madden, T. L., Matten, W. T., McGinnis, S. D., Merezhuk, Y., Raytselis, Y., Sayers, E. W., Tao, T., Ye, J., & Zaretskaya, I. (2013). BLAST: a more efficient report with usability improvements. Nucleic Acids Res., 41(Web Server issue), W29–33.

10. Brademan, D. R., Riley, N. M., Kwiecien, N. W., & Coon, J. J. (2019). Interactive peptide spectral annotator: A versatile web-based tool for proteomic applications. Mol. Cell. Proteom., 18(8, Supplement 1), S193–S201.

11. Buchan, D. W. A., & Jones, D. T. (2019). The PSIPRED protein analysis workbench: 20 years on. Nucleic Acids Research, 47(W1), W402–W407.

12. Chatterjee, A., Puri, S., Sharma, P. K., Deepa, P. R., & Chowdhury, S. (2023). Nature-inspired enzyme engineering and sustainable catalysis: biochemical clues from the world of plants and extremophiles [Review]. Front. Bioeng. Biotechnol., 11, 1229300.

13. Chen, P., Qi, F. X., Novak, J., Krull, R. E., & Caufield, P. W. (2001). Effect of amino acid substitutions in conserved residues in the leader peptide on biosynthesis of the lantibiotic mutacin II. FEMS Microbiol. Lett., 195(2), 139–144.

14. Chettri, D., Verma, A. K., Sarkar, L., & Verma, A. K. (2021). Role of extremophiles and their extremozymes in biorefinery process of lignocellulose degradation. Extremophiles, 25(3), 203–219.

15. Consortium, T. U. (2024). UniProt: the universal protein knowledgebase in 2025. Nucleic Acids Research, 53(D1), D609–D617.

16. De Leon, C. A., Levine, P. M., Craven, T. W., & Pratt, M. R. (2017). The sulfur-linked analogue of O-GlcNAc (S-GlcNAc) is an enzymatically stable and reasonable structural surrogate for O-GlcNAc at the peptide and protein levels. Biochemistry, 56(27), 3507–3517.

17. de Vos, W. M., Jung, G., & Sahl, H.-G. (1991). Appendix: definitions and nomenclature of lantibiotics. In G. Jung & H.-G. Sahl (Eds.), Nisin and Novel Lantibiotics (pp. 457–464). ESCOM.

18. Dhamdhere, G., & Zgurskaya, H. I. (2010). Metabolic shutdown in *Escherichia coli* cells lacking the outer membrane channel TolC. Mol. Microbiol., 77(3), 743–754.

19. Dirix, G., Monsieurs, P., Dombrecht, B., Daniels, R., Marchal, K., Vanderleyden, J., & Michiels, J. (2004). Peptide signal molecules and bacteriocins in Gram-negative bacteria: a genome-wide in silico screening for peptides containing a double-glycine leader sequence and their cognate transporters. Peptides, 25(9), 1425–1440.

20. Dirix, G., Monsieurs, P., Marchal, K., Vanderleyden, J., & Michiels, J. (2004). Screening genomes of Gram-positive bacteria for double-glycine-motif-containing peptides. Microbiology, 150(Pt 5), 1121–1126.

21. Dorenbos, R., Stein, T., Kabel, J., Bruand, C., Bolhuis, A., Bron, S., Quax, W. J., & van Dijl, J. M. (2002). Thiol-disulfide oxidoreductases are essential for the production of the lantibiotic sublancin 168. J. Biol. Chem., 277(19), 16682–16688.

22. Drummond, B. J., Loo, T. S., Patchett, M. L., & Norris, G. E. (2021). Optimised genetic tools allow the biosynthesis of glycocin F and analogues designed to test the roles of Gcc cluster genes in bacteriocin production. J. Bacteriol., 203(7), e00529–00520.

23. Eslami, S. M., & van der Donk, W. A. (2023). Proteases involved in leader peptide removal during RiPP biosynthesis. ACS Bio. Med. Chem. Au, 4(1), 20–36.

24. Fujinami, D., Garcia de Gonzalo, C. V., Biswas, S., Hao, Y., Wang, H., Garg, N., Lukk, T., Nair, S. K.., & van der Donk, W. A. (2021). Structural and mechanistic investigations of protein S-glycosyltransferases. Cell Chem. Biol., 28(12), 1740–1749.

25. Furgerson Ihnken, L. A., Chatterjee, C., & van der Donk, W. A. (2008). *In vitro* reconstitution and substrate specificity of a lantibiotic protease. Biochemistry, 47(28), 7352–7363.

26. Gai, J., File, M., Erdei, R., Czajlik, A., Marx, F., Galgóczy, L., Váradi, G., & Batta, G. (2025). Small Disulfide Proteins with Antifungal Impact: NMR Experimental Structures as Compared to Models of Alphafold Versions. Int. J. Mol. Sci., 26(3).

27. Garcia De Gonzalo, C. V., Denham, E. L., Mars, R. A., Stulke, J., van der Donk, W. A., & van Dijl, J. M. (2015). The phosphoenolpyruvate:sugar phosphotransferase system is involved in sensitivity to the glucosylated bacteriocin sublancin. Antimicrob. Agents Chemother., 59(11), 6844–6854.

28. Garcia De Gonzalo, C. V., Zhu, L., Oman, T. J., & van der Donk, W. A. (2014). NMR structure of the *S*-linked glycopeptide sublancin 168. ACS Chem. Biol., 9(3), 796–801.

29. Gerlt, J. A., Bouvier, J. T., Davidson, D. B., Imker, H. J., Sadkhin, B., Slater, D. R., & Whalen, K. L. (2015). Enzyme Function Initiative-Enzyme Similarity Tool (EFI-EST): A web tool for generating protein sequence similarity networks. Biochim. Biophys. Acta, 1854(8), 1019–1037.

30. Gomes, E., de Souza, A. R., Orjuela, G. L., Da Silva, R., de Oliveira, T. B., & Rodrigues, A. (2016). Applications and benefits of thermophilic microorganisms and their enzymes for industrial biotechnology. In M. Schmoll & C. Dattenböck (Eds.), Gene Expression Systems in Fungi: Advancements and Applications (pp. 459–492). Springer International Publishing. 10.1007/978-3-319-27951-0_21

31. Hart, G. W., Slawson, C., Ramirez-Correa, G., & Lagerlof, O. (2011). Cross Talk Between O-GlcNAcylation and Phosphorylation: Roles in Signaling, Transcription, and Chronic Disease. Annu. Rev. Biochem., 80, 825–858.

32. Hata, T., Tanaka, R., & Ohmomo, S. (2010). Isolation and characterization of plantaricin ASM1: a new bacteriocin produced by Lactobacillus plantarum A-1. Int. J. Food Microbiol., 137(1), 94–99.

33. Håvarstein, L. S., Diep, D. B., & Nes, I. F. (1995). A family of bacteriocin ABC transporters carry out proteolytic processing of their substrates concomitant with export. Mol. Microbiol., 16(2), 229–240.

34. Ishii, S., Yano, T., Ebihara, A., Okamoto, A., Manzoku, M., & Hayashi, H. (2010). Crystal structure of the peptidase domain of *Streptococcus* ComA, a bifunctional ATP-binding cassette transporter involved in the quorum-sensing pathway. J. Biol. Chem., 285(14), 10777–10785.

35. Ishii, S., Yano, T., & Hayashi, H. (2006). Expression and characterization of the peptidase domain of *Streptococcus pneumoniae* ComA, a bifunctional ATP-binding cassette transporter involved in quorum sensing pathway. J. Biol. Chem., 281(8), 4726–4731.

36. Izquierdo, E., Wagner, C., Marchioni, E., Aoude-Werner, D., & Ennahar, S. (2009). Enterocin 96, a novel class II bacteriocin produced by Enterococcus faecalis WHE 96, isolated from Munster cheese. Appl. Environ. Microbiol., 75(13), 4273–4276.

37. Kaunietis, A., Buivydas, A., Citavicius, D. J., & Kuipers, O. P. (2019). Heterologous biosynthesis and characterization of a glycocin from a thermophilic bacterium. Nat. Commun., 10(1), 1115.

38. Main, P., Hata, T., Loo, T. S., Man, P., Novak, P., Havlicek, V., Norris, G. E., & Patchett, M. L. (2020). Bacteriocin ASM1 is an O/S-diglycosylated, plasmid-encoded homologue of glycocin F. FEBS Lett., 594(7), 1196–1206.

39. Maky, M. A., Ishibashi, N., Nakayama, J., & Zendo, T. (2021). Characterization of the biosynthetic gene cluster of enterocin F4-9, a glycosylated bacteriocin. Microorganisms, 9(11), 2276.

40. Maky, M. A., Ishibashi, N., Zendo, T., Perez, R. H., Doud, J. R., Karmi, M., & Sonomoto, K. (2015). Enterocin F4-9, a novel O-linked glycosylated bacteriocin. Appl. Environ. Microbiol., 81(14), 4819–4826.

41. Maynard, J. C., Burlingame, A. L., & Medzihradszky, K. F. (2016). Cysteine S-linked N-acetylglucosamine (S-GlcNAcylation), A New Post-translational Modification in Mammals. Mol. Cell. Proteomics, 15(11), 3405–3411.

42. Montalbán-López, M., Scott, T. A., Ramesh, S., Rahman, I. R., van Heel, A. J., Viel, J. H., Bandarian, V., Dittmann, E., Genilloud, O., Goto, Y., Grande Burgos, M. J., Hill, C., Kim, S., Koehnke, J., Latham, J. A., Link, A. J., Martínez, B., Nair, S. K., Nicolet, Y., Rebuffat, S., Sahl, H.-G., Sareen, D., Schmidt, E. W., Schmitt, L., Severinov, K., Süssmuth, R. D., Truman, A. W., Wang, H., Weng, J.-K., van Wezel, G. P., Zhang, Q., Zhong, J., Piel, J., Mitchell, D. A., Kuipers, O. P., & van der Donk, W. A. (2021). New developments in RiPP discovery, enzymology and engineering. Nat. Prod. Rep., 38(1), 130–239.

43. Nagar, R., & Rao, A. (2017). An iterative glycosyltransferase EntS catalyzes transfer and extension of O– and S-linked monosaccharide in enterocin 96. Glycobiology, 27(8), 766–776.

44. Namboori, S. C., & Graham, D. E. (2008). Enzymatic analysis of uridine diphosphate N-acetyl-D-glucosamine. Anal. Biochem., 381(1), 94–100.

45. Nguyen, D. T., Mitchell, D. A., & van der Donk, W. A. (2024). Genome mining for new enzyme chemistry. ACS Catal., 14(7), 4536–4553.

46. Norris, G. E., & Patchett, M. L. (2016). The glycocins: in a class of their own. Curr. Opin. Struct. Biol., 40, 112–119.

47. Oberg, N., Zallot, R., & Gerlt, J. A. (2023). EFI-EST, EFI-GNT, and EFI-CGFP: Enzyme function initiative (EFI) web resource for genomic enzymology tools. J. Mol. Biol., 435(14), 168018.

48. Oman, T. J., Boettcher, J. M., Wang, H., Okalibe, X. N., & van der Donk, W. A. (2011). Sublancin is not a lantibiotic but an *S*-linked glycopeptide. Nat. Chem. Biol., 7(2), 78–80.

49. Oman, T. J., & van der Donk, W. A. (2010). Follow the leader: the use of leader peptides to guide natural product biosynthesis. Nat. Chem. Biol., 6(1), 9–18.

50. Palaniappan, K., Chen, I. A., Chu, K., Ratner, A., Seshadri, R., Kyrpides, N. C., Ivanova, N. N., & Mouncey, N. J. (2020). IMG-ABC v.5.0: an update to the IMG/Atlas of Biosynthetic Gene Clusters Knowledgebase. Nucleic Acids Res., 48(D1), D422–d430.

51. Pan, A., Liu, X., Han, H., Gao, S. Q., & Lin, Y. W. (2025). Disruption of a potential disulfide bond of Cys65-Cys141 on the structure and stability of globin X from zebrafish. Phys. Chem. Chem. Phys., 27(5), 2828–2833.

52. Pei, J., Pang, Q., Zhao, L., Fan, S., & Shi, H. (2012). *Thermoanaerobacterium thermosaccharolyticum* β-glucosidase: a glucose-tolerant enzyme with high specific activity for cellobiose. Biotechnol. Biofuels Bioprod., 5(1), 31.

53. Ren, H., Biswas, S., Ho, S., van der Donk, W. A., & Zhao, H. (2018). Rapid discovery of glycocins through pathway refactoring in *Escherichia coli*. ACS Chem. Biol., 13(10), 2966–2972.

54. Repka, L. M., Chekan, J. R., Nair, S. K., & van der Donk, W. A. (2017). Mechanistic understanding of lanthipeptide biosynthetic enzymes. Chem. Rev., 117, 5457–5520.

55. Shannon, P., Markiel, A., Ozier, O., Baliga, N. S., Wang, J. T., Ramage, D., Amin, N., Schwikowski, B., & Ideker, T. (2003). Cytoscape: a software environment for integrated models of biomolecular interaction networks. Genome Res., 13(11), 2498–2504.

56. Sharma, Y., Ahlawat, S., & Rao, A. (2021). Biochemical characterization of an inverting S/O-HexNAc-transferase and evidence of S-linked glycosylation in Actinobacteria. Glycobiology, 32(2), 148–161.

57. Singh, V., & Rao, A. (2021). Distribution and diversity of glycocin biosynthesis gene clusters beyond Firmicutes. Glycobiology, 31(2), 89–102.

58. Stepper, J., Shastri, S., Loo, T. S., Preston, J. C., Novak, P., Man, P., Moore, C. H., Havlicek, V., Patchett, M. L., & Norris, G. E. (2011). Cysteine S-glycosylation, a new post-translational modification found in glycopeptide bacteriocins. FEBS Lett., 585, 645–650.

59. Venugopal, H., Edwards, P. J., Schwalbe, M., Claridge, J. K., Libich, D. S., Stepper, J., Loo, T., Patchett, M. L., Norris, G. E., & Pascal, S. M. (2011). Structural, dynamic, and chemical characterization of a novel S-glycosylated bacteriocin. Biochemistry, 50(14), 2748–2755.

60. Wang, H., Han, Y., Wang, X., Jia, Y., Zhang, Y., Müller, R., & Huo, L. (2023). Genome mining of myxopeptins reveals a class of lanthipeptide-derived linear dehydroamino acid-containing peptides from Myxococcus sp. MCy9171. ACS Chem. Biol., 18(10), 2163–2169.

61. Wang, H., Oman, T. J., Zhang, R., Garcia De Gonzalo, C. V., Zhang, Q., & van der Donk, W. A. (2014). The glycosyltransferase involved in thurandacin biosynthesis catalyzes both O– and S-glycosylation. J. Am. Chem. Soc., 136(1), 84–87.

62. Wang, H., & van der Donk, W. A. (2011). Substrate selectivity of the sublancin S-glycosyltransferase. J. Am. Chem. Soc., 133(41), 16394–16397.

63. Wang, H., Zhao, X., Li, D., Meng, L., Liu, S., Zhang, Y., & Huo, L. (2025). Marine metagenome mining reveals lanthipeptides colwesin A–C, exhibiting novel ring topology and anti-inflammatory activity. ACS Synth. Biol.

64. Wang, J., Ge, X., Zhang, L., Teng, K., & Zhong, J. (2016). One-pot synthesis of class II lanthipeptide bovicin HJ50 via an engineered lanthipeptide synthetase. Sci. Rep., 6, 38630.

65. Wang, X., Chen, X., Wang, Z. J., Zhuang, M., Zhong, L., Fu, C., Garcia, R., Müller, R., Zhang, Y., Yan, J., Wu, D., & Huo, L. (2023). Discovery and characterization of a myxobacterial lanthipeptide with unique biosynthetic features and anti-inflammatory activity. J. Am. Chem. Soc., 145(30), 16924–16937.

66. Wu, C., Biswas, S., Garcia De Gonzalo, C. V., & van der Donk, W. A. (2019). Investigations into the mechanism of action of sublancin. ACS Infect. Dis., 5(3), 454–459.

67. Wu, K. H., & Tai, P. C. (2004). Cys32 and His105 are the critical residues for the calcium-dependent cysteine proteolytic activity of CvaB, an ATP-binding cassette transporter. J. Biol. Chem., 279(2), 901–909.

68. Zhu, D., Adebisi, W. A., Ahmad, F., Sethupathy, S., Danso, B., & Sun, J. (2020). Recent development of extremophilic bacteria and their application in biorefinery [Review]. Front. Bioeng. Biotechnol., 8, 483.

