## Supplemental figures for "Characterization of a novel glycocin from a thermophile"

TABLE S1. BACTERIA TESTED FOR ANTIMICROBIAL ACTIVITY WITH THERMOGLYCOCIN 15

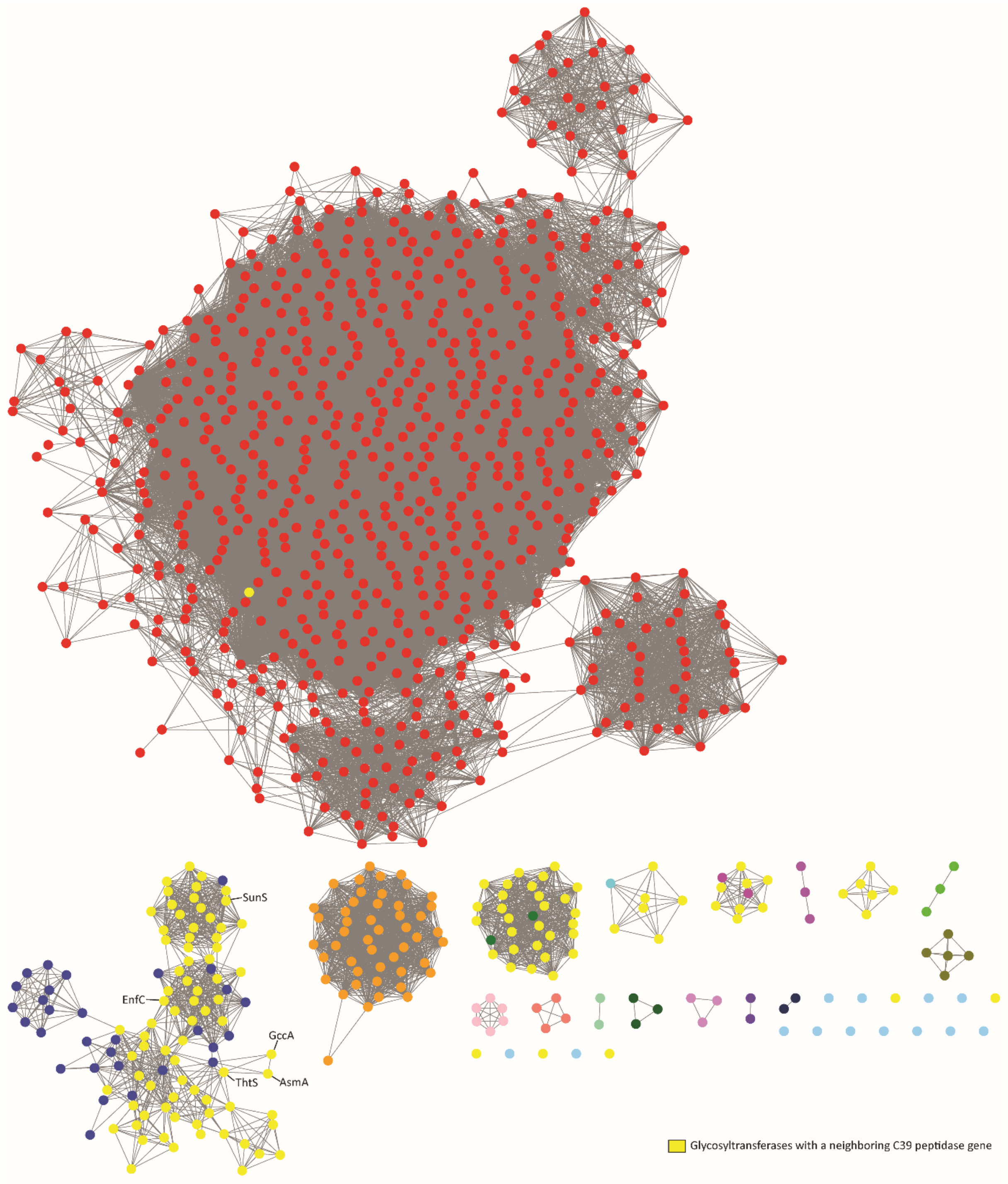

Fig. S1. Sequence similarity network of ThtS related proteins.

Sequence similarity network (alignment threshold of 50) of proteins related to ThtS generated using the UniProt database (Consortium, 2024). Nodes highlighted in yellow show proteins with a neighboring (within 10 genes) C39 peptidase/ABC transporter determined by gene neighborhood network analysis (Oberg et al., 2023). Known glycosyltransferases involved in glycocin biosynthesis are labeled, including AsmA (Main et al., 2020) and GccA (Stepper et al., 2011), first neighbors of ThtS, and EnfC (Maky et al., 2015), and SunS (Oman et al., 2011).

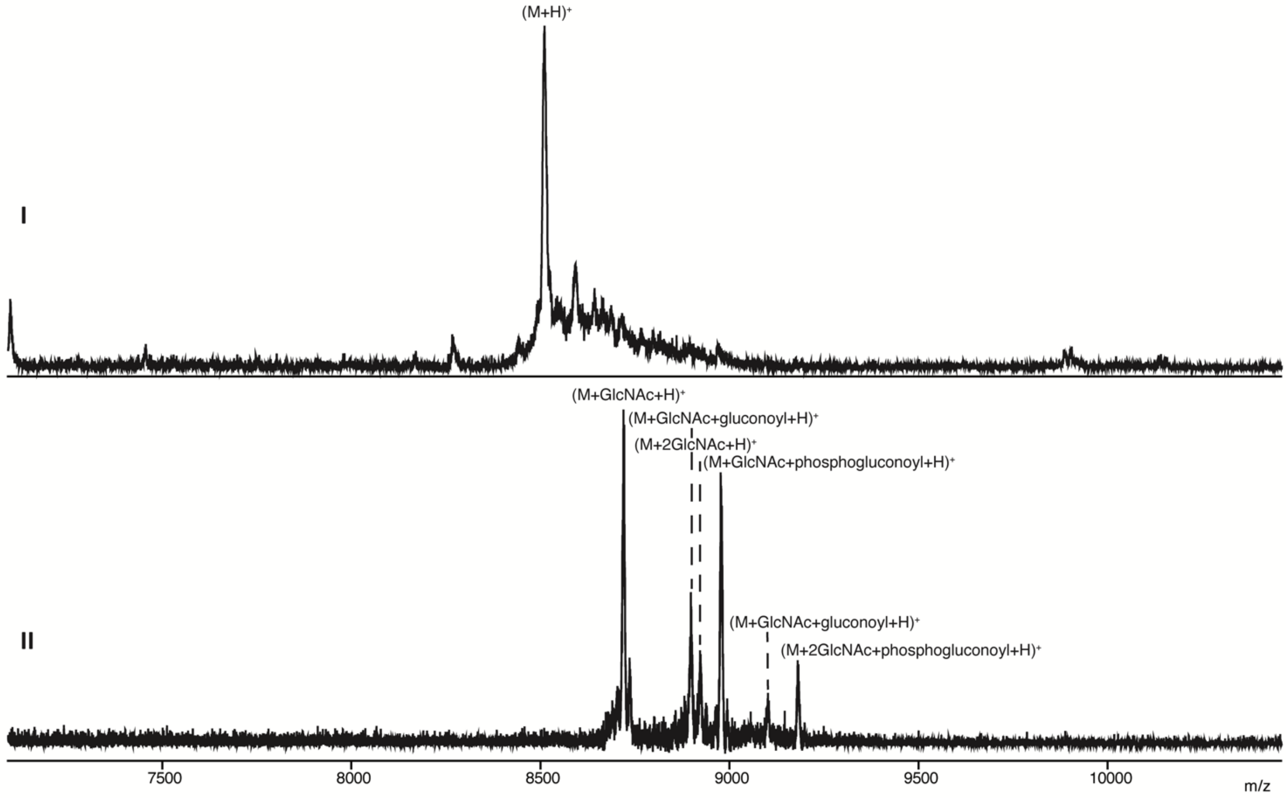

Fig. S2. His$_6$-ThtA modification by ThtS in *E. coli*.

(I) MALDI-TOF mass spectrum of His$_6$-ThtA with three disulfide bonds purified from *E. coli* expression. $(M+H)^+$ calculated *m/z*: 8501.7, observed *m/z*: 8503.6. (II) MALDI-TOF mass spectrum of His$_6$-ThtA (disulfide bonds reduced with TCEP) purified after co-expression with ThtS. Additional modifications are due to gluconoylation and phosphogluconoylation of the His-tag (Geoghegan et al., 1999). $(M + GlcNAc+H)^+$ calculated *m/z*: 8710.8, observed *m/z*: 8714.8; $(M + GlcNAc + gluconoyl+H)^+$ calculated *m/z*: 8888.8, observed *m/z*: 8892.5; $(M + 2GlcNAc+H)^+$ calculated *m/z*: 8913.9, observed *m/z*: 8921.2; $(M + GlcNAc + phosphogluconoyl+H)^+$ calculated *m/z*: 8968.8, observed *m/z*: 8973.2; $(M + 2GlcNAc + gluconoyl+H)^+$ calculated *m/z*: 9091.9, observed *m/z*: 9101.9; $(M + 2GlcNAc + phosphogluconoyl+H)^+$ calculated *m/z*: 9171.9, observed *m/z*: 9180.7

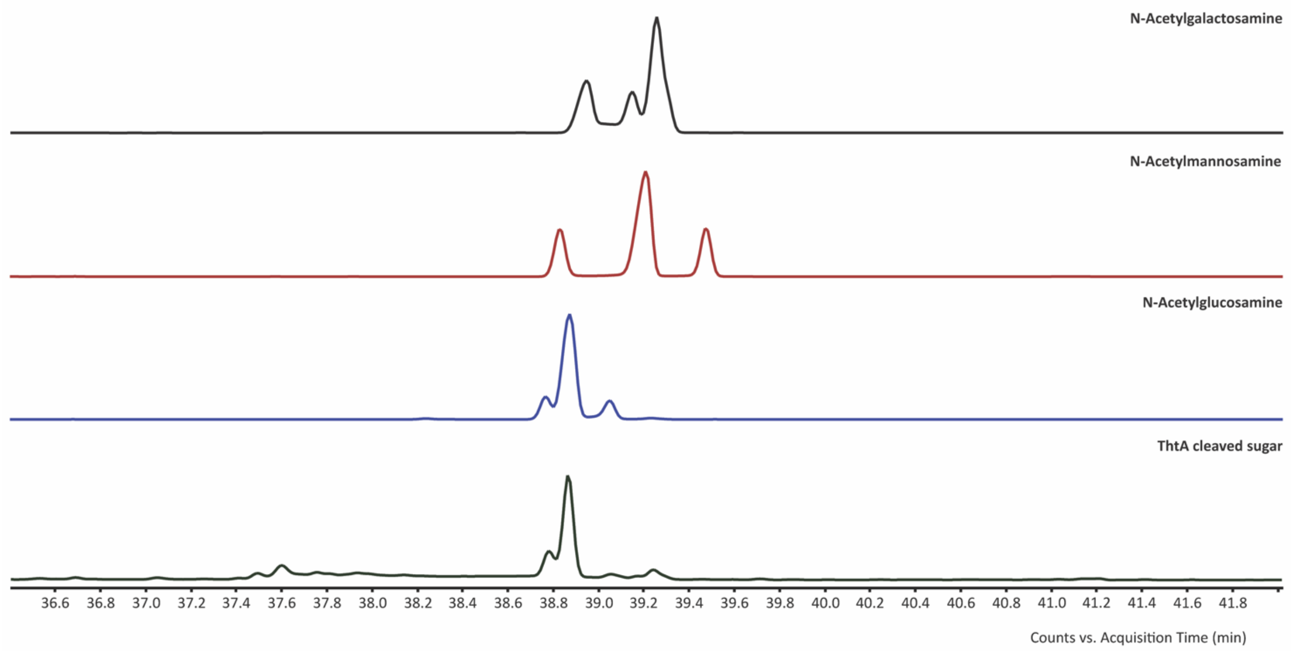

Fig. S3. Identification of ThtA glycosylation using GC-MS.

GC-MS analysis of the sugar cleaved from ThtA after derivatization as explained in the methods section (bottom) in comparison to N-acetylated hexose standards that were derivatized similarly.

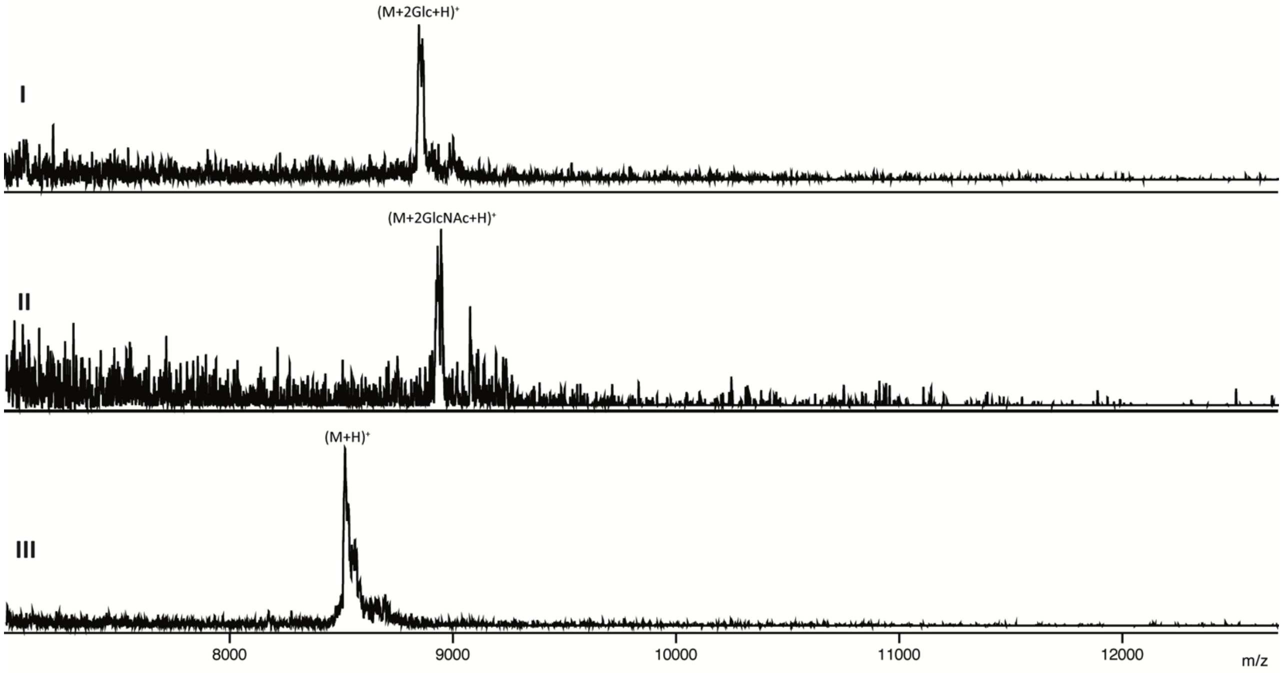

Fig. S4. *In vitro* glucosylation of ThtA by ThtS.

MALDI-TOF mass spectra of $His_6$-ThtA incubated with $His_6$-ThtS and UDP-glucose or UDP-GlcNAc. (I) (M + 2 Glc +H)$^+$ calculated *m/z*: 8831.8, observed *m/z*: 8846.9; (II) (M+2GlcNAc+H)$^+$ calculated *m/z*: 8913.9, observed *m/z*:8930.9; and (III) unmodified $His_6$-ThtA (M+H)$^+$ calculated *m/z*: 8507.6971, observed *m/z*:. 8516.768

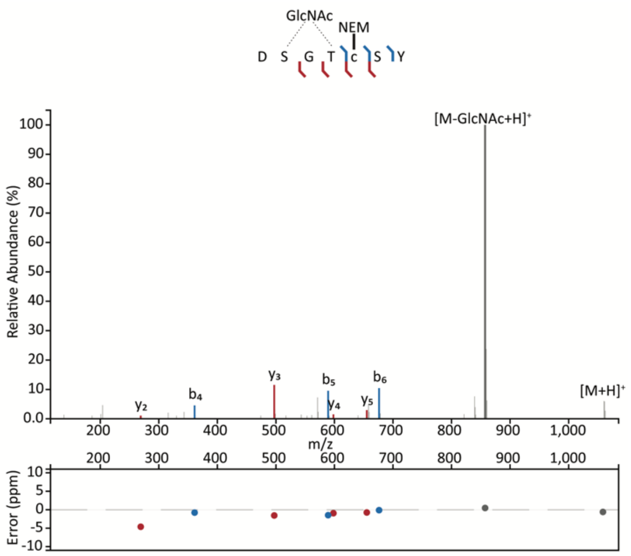

Fig. S5. Chymotrypsin digest fragment of mThtA.

ESI-LC-MS/MS spectrum of mThtA fragment encompassing residues 19 to 25 $(M+H)^+$ calculated *m/z*: 1,060.3776, observed *m/z*: 1060.3773; $(M - GlcNAc+H)^+$ calculated *m/z*: 857.2982, observed *m/z*: 857.2986. mThtA was alkylated with NEM and digested by chymotrypsin. Fragmentation data shows the GlcNAc is cleaved from the peptide during fragmentation, before the peptide bonds are broken. All fragments lack GlcNAc. Figure prepared using the Interactive Peptide Annotator Webtool (Brademan et al., 2019).

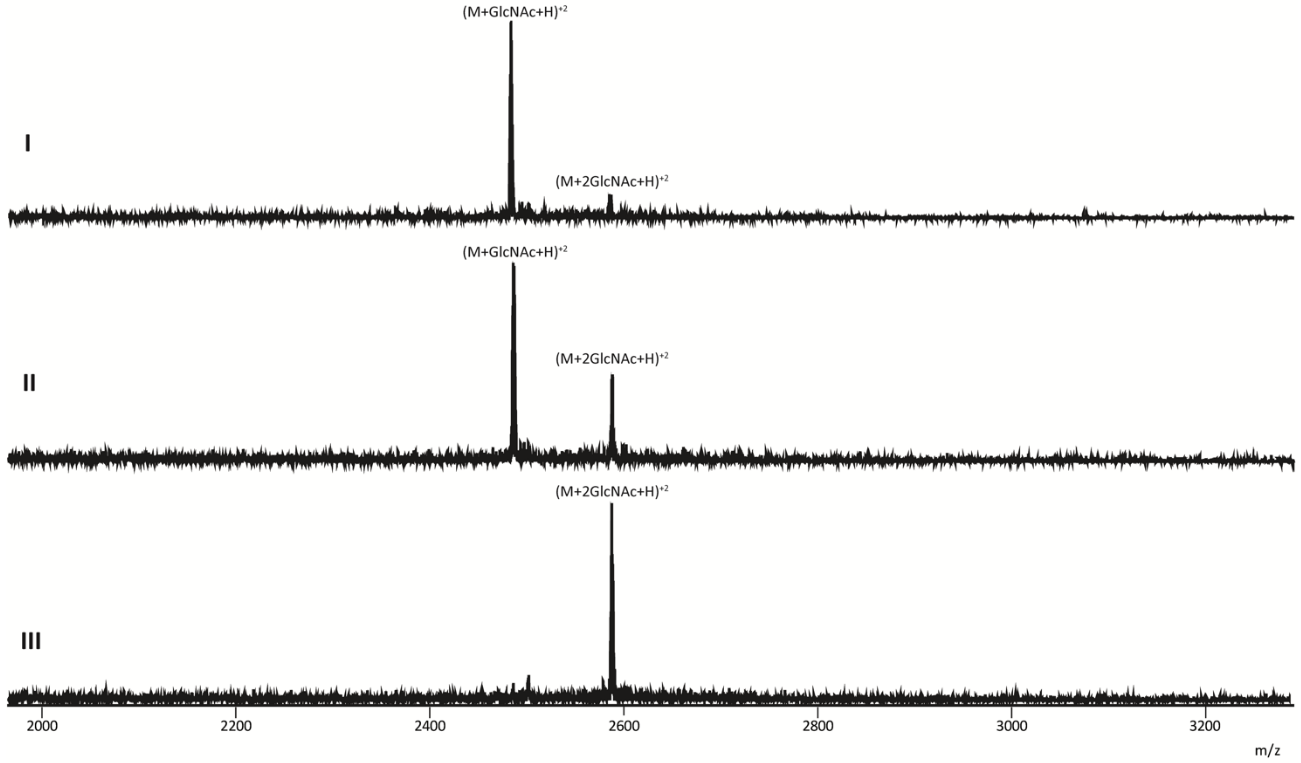

Fig. S6. GlcNAcylation of mThtA core peptide by ThtS.

MALDI-TOF MS of doubly charged $His_6$-ThtA co-expressed with ThtS in *E. coli* and purified as a mixture of mono- and di-GlcNAcylated peptides followed by digestion with LysC, which removes the putative leader peptide after Lys-1 (see Fig. 1 main text). (I) Monoglycosylated peptide: calculated *m/z*: 2484.5, observed *m/z*: 2483.1. (II) Incubation with ThtS and UDP-GlcNAc *in vitro* shows conversion to di-GlcNAcylated peptide after 1 h. Calculated *m/z*: 2586.1, observed *m/z*: 2587.0. (III) Incubation with ThtS and UDP-GlcNAc after 16 h. Calculated *m/z*: 2586.1, observed *m/z*: 2586.1).

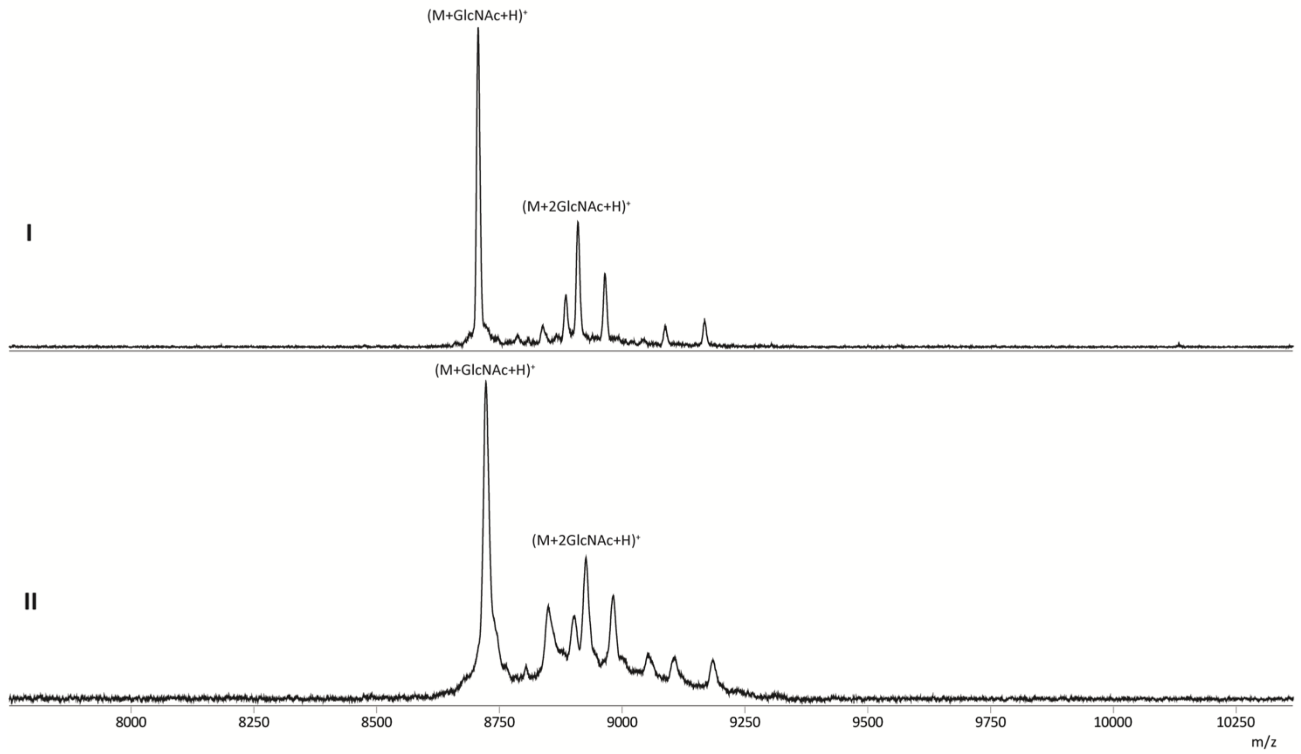

Fig. S7. Determination of the number of free cysteines in mThtA.

$His_6$-ThtA co-expressed with ThtS and purified from *E. coli* as a mixture of mono- and diGlcNAcylated peptide. MALDI-TOF mass spectra of (I) (M + GlcNAc+H)$^+$ calculated *m/z*: 8704.7, observed *m/z*: 8701.0; (M + 2 GlcNAc +H)$^+$ calculated *m/z*: 8907.8, observed *m/z*: 8905.4. (II) The peptide was incubated with NEM in the absence of reductant to determine the number of free cysteine residues. No adducts were observed, showing full formation of three disulfides. (M + GlcNAc+H)$^+$ calculated *m/z*: 8704.7, observed *m/z*: 8713.0 (M + 2 GlcNAc+H)$^+$ calculated *m/z*: 8907.8, observed *m/z*: 8919.5.

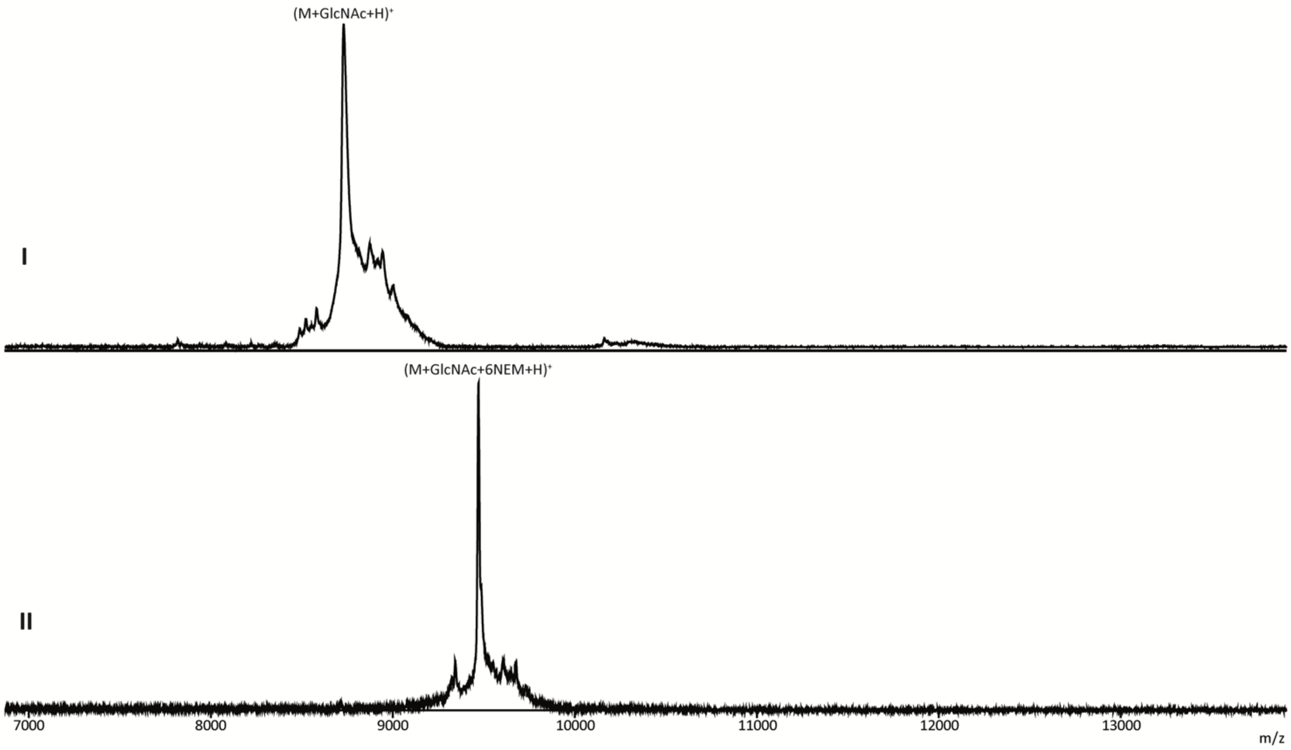

Fig. S8. Determination of the number of cysteines involved in disulfide bonding.
Alkylation of monoGlcNAcylated His$_6$-ThtA, from co-expression with ThtS, to determine the number of cysteine residues involved in disulfide bonding. MALDI-TOF mass spectra of (I) His$_6$-ThtA + GlcNAc reduced with TCEP. $(M + GlcNAc+H)^+$ calculated *m/z*: 8710.8, observed *m/z*: 8712.9, and (II) His$_6$-ThtA + GlcNAc reduced with TCEP and incubated with NEM. $(M + GlcNAc + 6\ NEM+H)^+$ calculated *m/z*: 9461.1, observed *m/z*: 9461.3.

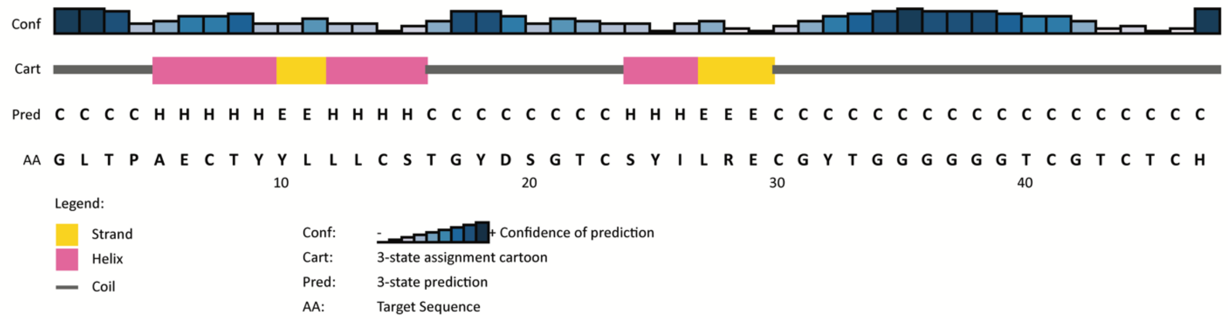

Fig. S9. Prediction of ThtA secondary structure.

PSIPRED prediction of the ThtA core peptide secondary structure, showing two α-helices connected by a coil.

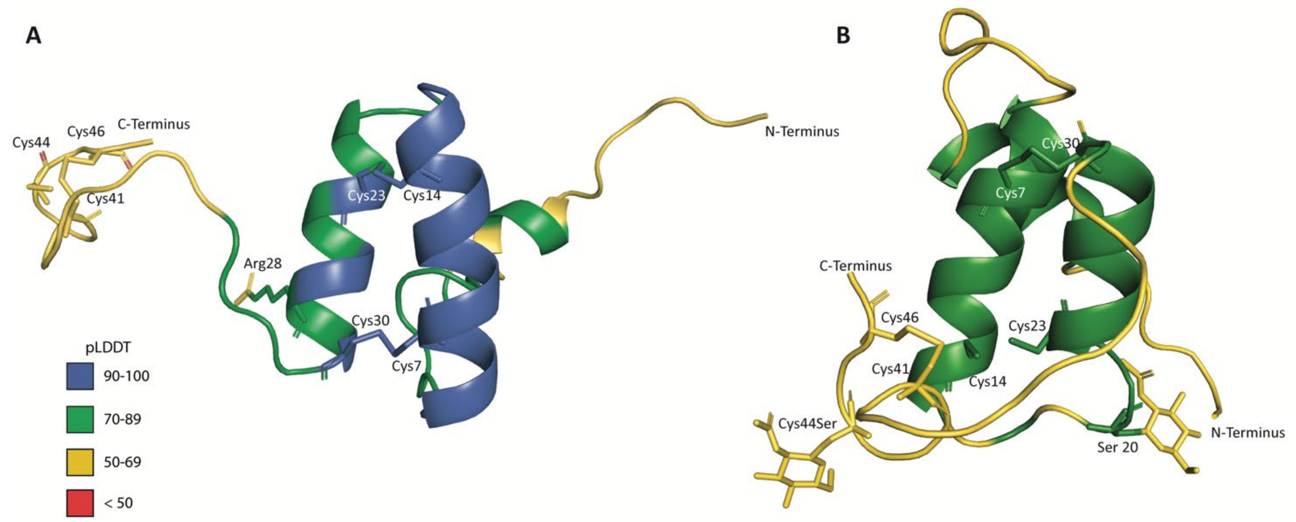

Fig. S10. AlphaFold3 models of thermoglycocin.

AlphaFold3 model of A) Thermoglycocin showing disulfide bonds predicted between Cys7 and Cys30, Cys14 and Cys23, and Cys41 and Cys46. B) Thermoglycocin with Cys44 mutated to Ser to allow for glycosylation on residues 20 and 44 using AlphaFold3. The peptide is colored by pLDDT confidence values from low (red) to high (blue).

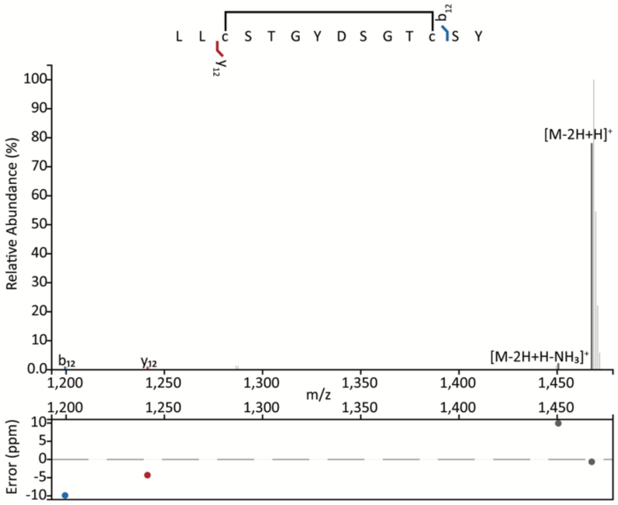

Fig. S11. LC MS/MS confirmation of a disulfide bond between Cys14 and Cys23.

LC-MS/MS of thermolysin-digested monoglycosylated ThtA fragment containing residues 12-25 with a disulfide bond connecting Cys14 and Cys23, $(M\text{-}2H+H)^+$ calculated *m/z*: 1467.5767, observed *m/z*: 1467.5757. Figure prepared using the Interactive Peptide Annotator Webtool (Brademan et al., 2019).

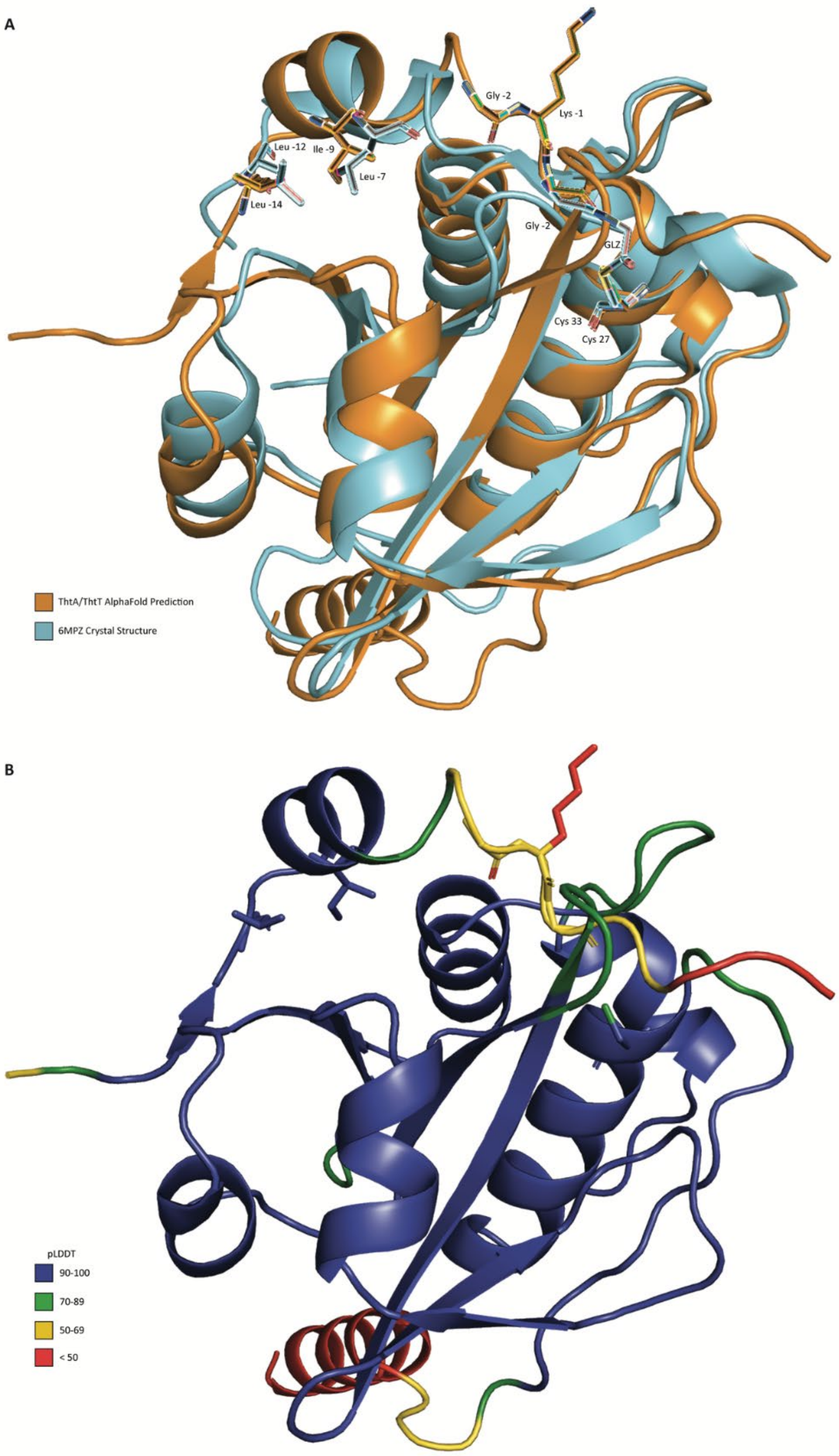

Fig. S12. Overlay of ThtA/ThtS predicted structure with LahA/LahT crystal structure.

Overlay of the LahT crystal structure with a bound inhibitor (GLZ) based on the LahA sequence (PDB ID: 6MPZ) with the AlphaFold prediction of ThtT and ThtA leader peptide structure. Highlighted residues show that the hydrophobic residues of the ThtA leader peptide at positions −9 and −14 overlap with the hydrophobic residues of the LahA leader peptide at positions −7 and −12. The GG motif of LahA (shown here as Gly and covalently bound amino-acetaldehyde) binds in a similar fashion as the GK motif in ThtA.

Table S1. Bacteria tested for antimicrobial activity with thermoglycocin using agar diffusion assays.

| **Strain tested** | **Bioactivity** |
|---|---|
| *B. subtilis* 168 | NOT OBSERVED |
| *B. subtilis* 168 ΔSPβ | NOT OBSERVED |
| *E. coli* BL21 | NOT OBSERVED |
| *L. lactis* | NOT OBSERVED |
| *M. luteus* | NOT OBSERVED |
| *E. faecalis* | NOT OBSERVED |
| *B. cereus* strain TZ417 | NOT OBSERVED |
| *G. thermodenitrificans* | NOT OBSERVED |
| *B. megaterium* | NOT OBSERVED |
| *E. faecium* | NOT OBSERVED |
| *S. aureus* | NOT OBSERVED |
| *K. pneumoniae* | NOT OBSERVED |
| *A. baumannii* | NOT OBSERVED |
| *P. aeruginosa* | NOT OBSERVED |
| *E. cloacae* | NOT OBSERVED |

### References

Brademan, D. R., Riley, N. M., Kwiecien, N. W., & Coon, J. J. (2019). Interactive peptide spectral annotator: A versatile web-based tool for proteomic applications. *Mol. Cell. Proteom.*, *18*(8, Supplement 1), S193-S201. https://doi.org/10.1074/mcp.TIR118.001209

Consortium, T. U. (2024). UniProt: the universal protein knowledgebase in 2025. *Nucleic Acids Research*, *53*(D1), D609-D617. https://doi.org/10.1093/nar/gkae1010

Geoghegan, K. F., Dixon, H. B., Rosner, P. J., Hoth, L. R., Lanzetti, A. J., Borzilleri, K. A., Marr, E. S., Pezzullo, L. H., Martin, L. B., LeMotte, P. K., McColl, A. S., Kamath, A. V., & Stroh, J. G. (1999). Spontaneous alpha-N-6-phosphogluconoylation of a "His tag" in *Escherichia coli*: the cause of extra mass of 258 or 178 Da in fusion proteins. *Anal. Biochem.*, *267*(1), 169-184. https://doi.org/10.1006/abio.1998.2990

Main, P., Hata, T., Loo, T. S., Man, P., Novak, P., Havlicek, V., Norris, G. E., & Patchett, M. L. (2020). Bacteriocin ASM1 is an O/S-diglycosylated, plasmid-encoded homologue of glycocin F. *FEBS Lett.*, *594*(7), 1196-1206. https://doi.org/10.1002/1873-3468.13708

Maky, M. A., Ishibashi, N., Zendo, T., Perez, R. H., Doud, J. R., Karmi, M., & Sonomoto, K. (2015). Enterocin F4-9, a novel O-linked glycosylated bacteriocin. *Appl. Environ. Microbiol.*, *81*(14), 4819-4826. https://doi.org/10.1128/aem.00940-15

Oberg, N., Zallot, R., & Gerlt, J. A. (2023). EFI-EST, EFI-GNT, and EFI-CGFP: Enzyme Function Initiative (EFI) web resource for genomic enzymology tools. *J. Mol. Biol.*, *435*(14), 168018. https://doi.org/10.1016/j.jmb.2023.168018

Oman, T. J., Boettcher, J. M., Wang, H., Okalibe, X. N., & van der Donk, W. A. (2011). Sublancin is not a lantibiotic but an *S*-linked glycopeptide. *Nat. Chem. Biol.*, *7*(2), 78-80.

Stepper, J., Shastri, S., Loo, T. S., Preston, J. C., Novak, P., Man, P., Moore, C. H., Havlicek, V., Patchett, M. L., & Norris, G. E. (2011). Cysteine S-glycosylation, a new post-translational modification found in glycopeptide bacteriocins. *FEBS Lett.*, *585*, 645-650.
